# High-density extracellular probes reveal dendritic backpropagation and facilitate neuron classification

**DOI:** 10.1101/376863

**Authors:** Xiaoxuan Jia, Josh Siegle, Corbett Bennett, Sam Gale, Daniel R Denman, Christof Koch, Shawn R. Olsen

## Abstract

Different neuron types serve distinct roles in neural processing. Extracellular electrical recordings are extensively used to study brain function but are typically blind to cell identity. Morpho-electric properties of neurons measured on spatially dense electrode arrays might be useful for distinguishing neuron types. Here we used Neuropixels probes to record from cortical and subcortical regions of the mouse brain. Extracellular waveforms of each neuron were detected across many channels and showed distinct spatiotemporal profiles among brain regions. Classification of neurons by brain region was improved with multi-channel compared to single-channel waveforms. In visual cortex, waveform clustering identified the canonical regular spiking (RS) and fast spiking (FS) classes, but also uncovered a subclass of RS units with unidirectional backpropagating action potentials (BAPs). Moreover, BAPs were observed in many hippocampal RS cells. Overall, waveform analysis of spikes from high-density probes aids neuron identification and can reveal dendritic backpropagation.

## INTRODUCTION

Brain networks are composed of diverse cell types that play distinct roles in neural dynamics and processing. For example, in the neocortex, excitatory pyramidal neurons provide local recurrent processing and send long-range projections for information propagation (Harris and Shepherd, 2015; Spruston, 2008), while inhibitory neurons perform gain modulation, control spike timing and rhythms, and shape receptive field properties (Isaacson and Scanziani, 2011; Kepecs and Fishell, 2014; Markram et al., 2004). Thus, a mechanistic understanding of brain function requires a cell type-specific approach. Neuronal cell types are defined by various properties including gene expression, morphology, physiology, and connectivity (Baden et al., 2016; Gouwens et al., 2018; Harris and Shepherd, 2015; Kim et al., 2017; Markram et al., 2004; Tasic et al., 2016; Zeng and Sanes, 2017). While imaging of genetically-encoded calcium sensors can be used to measure activity of identified neuronal subpopulations (Luo et al., 2008), this method has much lower temporal resolution compared to electrophysiological recordings and can be difficult in deep brain structures. A technique called optotagging can link extracellular spike measurements to cell types by directly photo-stimulating cells that express a light-sensitive opsin under genetic control (Cohen et al., 2012; Kvitsiani et al., 2013; Lima et al., 2009), but this is largely restricted to transgenic systems and typically only labels one cell population per experiment.

Extracellular electrical recordings are widely used to measure single neuron spiking activity in vivo during active behavior. Previous studies have shown that action potential shape can provide information about the cell type being recorded (Kawaguchi, 1993; McCormick et al., 1985). Conventionally, single unit waveforms are divided into two broad categories: regular spiking (RS), which represent pyramidal neurons and some inhibitory neurons, and fast spiking neurons (FS), which largely correspond to inhibitory interneurons (Andermann et al., 2004; Bortone et al., 2014; Bruno and Simons, 2002; Mitchell et al., 2007; Niell and Stryker, 2008; Peyrache et al., 2012; Sirota et al., 2008; Swadlow, 2003). In general, RS neurons are characterized by broader action potentials with spike frequency adaptation, while FS neurons have relatively brief duration action potentials with little adaptation (Hu et al., 2014; Markram et al., 2004). This classification is supported by the correlation between extracellular waveform and intrinsic electrophysiological properties (Anastassiou et al., 2015; Buzsáki et al., 2012; Gold et al., 2006; Henze et al., 2000). The ability to more generally link cell classes to extracellular action potential waveform features would enhance many studies of circuit-level functions in the brain.

High-density silicon probe technology provides increased spatial sampling of extracellular waveforms from single units in vivo (Jun et al., 2017; Neto et al., 2016; Shobe et al., 2015). The density of electrode contacts on these probes allows each unit to be visible on many sites simultaneously, thereby providing a rich spatiotemporal extracellular waveform profile for each unit. Here we used Neuropixels probes to make large-scale electrophysiological recordings in the awake mouse brain and performed analysis to investigate spike waveform features that might facilitate cell classification based on extracellular measurements (Figure 1A, B). Since morpho-electric properties are key attributes of cell types (Ascoli et al., 2007; Gouwens et al., 2018; Jiang et al., 2015; Zeng and Sanes, 2017), we hypothesized that the spatial dimension of the extracellular electrical field captured with Neuropixels could improve cell classification (Figure 1B). For example, pyramidal neurons with apical dendrites can show backpropagating action potentials (Bereshpolova et al., 2007; Buzsáki and Kandel, 1998; Stuart and Sakmann, 1994; Stuart et al., 1997; Waters et al., 2005), and these might be visible as traveling waves along the probe which could be used to differentiate pyramidal neurons from cells with distinct dendritic backpropagation properties and/or morphologies.

We first sought to test whether multi-channel waveforms can facilitate cell identification across different brain areas, since morpho-electric properties show distinct variation across cortical and subcortical structures (Ascoli et al., 2007; Bean, 2007; Stuart et al., 1997). We applied classification and clustering algorithms to single and multi-channel features of extracellular spike waveforms recorded in the visual cortex, hippocampus, thalamus, superior colliculus, and cerebellum of the awake mouse brain (Figure 1C, D). By combining signals from multiple channels, we could better classify cells from different brain regions. Within the local circuitry of the neocortex and hippocampus, our clustering recapitulated the conventional RS and FS division but suggested additional waveform diversity. We validated the FS cluster by linking to genetic cell types in the visual cortex by optotagging PV+ neurons. A substantial number of RS units in both visual cortex and hippocampus showed evidence of backpropagating action potentials (BAPs), but the FS cluster did not. These putative BAPs are reliably observed in vivo, and their signature can help differentiate cells within local circuits of the cortex and hippocampus. We conclude that dense sampling of the extracellular waveform with next-generation extracellular probes provides additional information for cell type and brain region classification based purely on spike waveform, which can complement additional methodologies for dissecting cell type-specific neural network functions.

**Figure 1.**
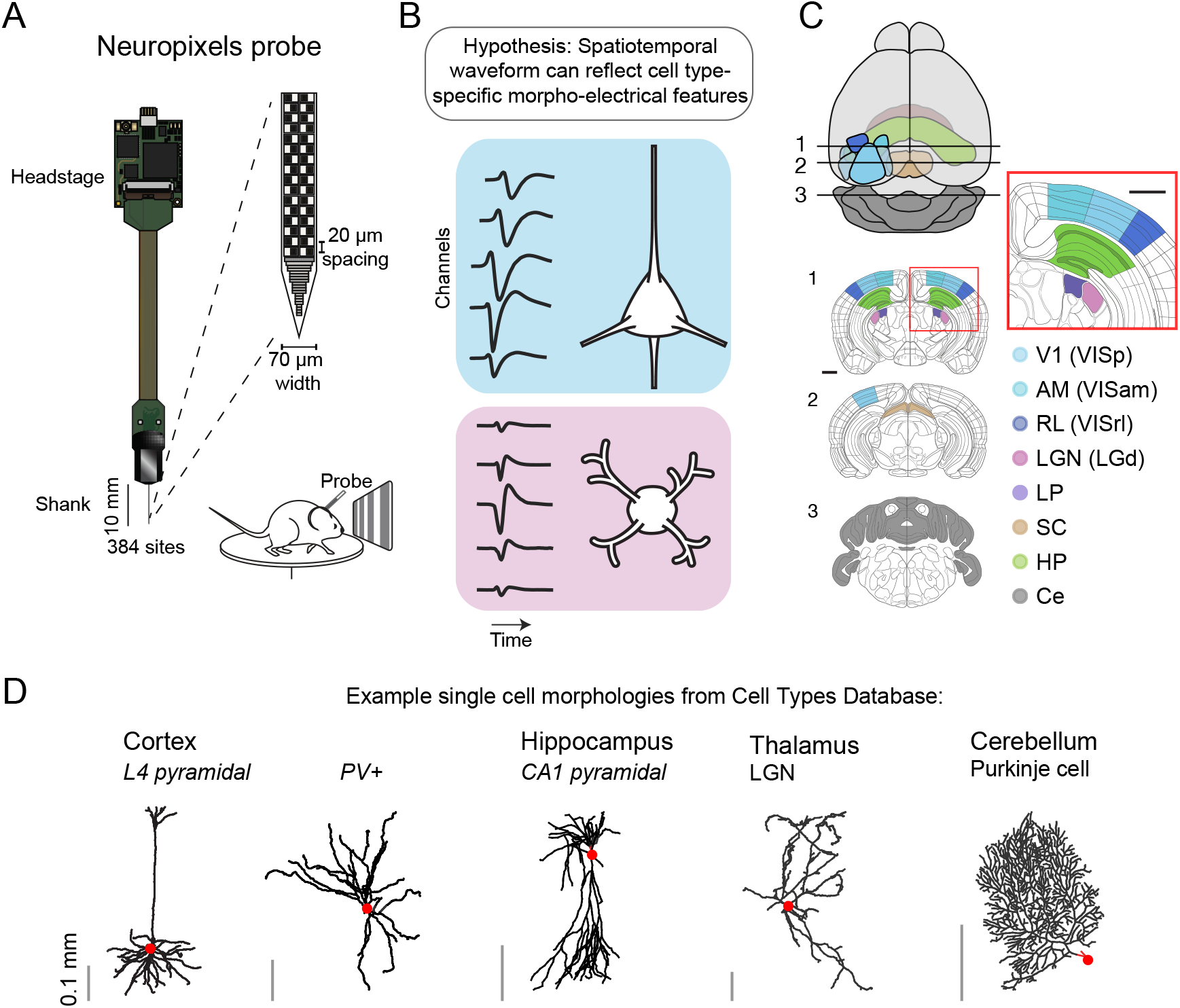
Experimental setup and hypothesis. **A)** Data are collected with Neuropixels probes inserted into the awake mouse brain. The 384 electrode sites are densely arranged along the linear shank of the silicon probe (20 um vertical spacing, 2 sites per row). Black squares indicate the location of recording sites. **B)** Schematic illustrating our hypothesis that extracellular waveforms at different spatial locations relative to the neuron can reflect cell type-specific morpho-electric features. **C)** Illustrations of brain regions targeted for recordings (from Allen Mouse Brain Atlas). In the visual cortex, we recorded from V1, and two higher visual areas AM and RL (different blues). In subcortical regions, we recorded from dorsal hippocampus (HP; green), LGN (pink), LP (purple), superior colliculus (SC; brown), and cerebellum (Ce; gray). **D)** Example cell reconstructions from different brain regions to illustrate morphological diversity. Reconstructions are from Allen Cell Types Database (http://celltypes.brain-map.org/) and NeuroMorpho Database (http://www.neuromorpho.org/). Dendrites are shown in black and cell body location is denoted with red circle.

## RESULTS

We analyzed extracellular action potentials recorded from 43 adult mice using Neuropixels probes. In each recording session, the mouse was awake and head-fixed on a wheel to allow free running behavior; in some experiments mice were also exposed to visual stimuli (Figure 1A). After spike sorting with semi-automated algorithms (Pachitariu et al., 2016; Rossant et al., 2016a), we recovered 2818 well-isolated single units with signal-to-noise ratio larger than 1.5. These units were recorded from 8 brain regions: primary visual cortex V1 (VISp, 1111 units), cortical visual area AM (VISam, 234 units), cortical visual area RL (VISrl, 264 units), hippocampus (HP, 369 units; mostly dorsal CA1, see Methods), lateral geniculate nucleus (LGN, 106 units), lateral posterior nucleus (LP, 485 units), superior colliculus (SC, 171 units), and cerebellum (Ce, 78 units) (Figure 2A). To verify the brain regions recorded by each probe, we used post hoc histology, imaging, and annotation using the Allen Mouse Common Coordinate Framework (Supplementary Figure 1). Subregions of visual cortex (VISp, VISam, VISrl) were determined by functional retinotopic mapping prior to the experiment and these maps were used to guide probe insertion. Thus, for each recording we could label each sorted unit with its brain region. The action potential waveforms analyzed in this paper refer to the mean waveform for each sorted unit, which is calculated by taking a bootstrapped average (number_of_spikes=100; number_of_repetitions=100) from all spikes aligned by their trough (see Methods). We define the 1-channel waveform for a given unit as the mean action potential recorded on the channel with the largest amplitude—often this channel is assumed to be closest to the soma (Buzsáki and Kandel, 1998), and we follow that convention here.

### Comparison of single channel waveforms from different brain regions

We first investigated whether neurons in different brain regions have distinct single-channel (1-channel) waveform profiles (Figure 2A; each line is the mean waveform for one unit normalized by amplitude). Units in the visual cortex and hippocampus show a diversity of 1-channel waveform shapes including both narrow and wide spikes; in contrast, 1-channel waveforms from the cerebellum are more consistently narrow, while thalamic cells are more consistently broad (Figure 2A). To quantitatively compare spikes, we extracted a series of features from the 1-channel waveforms including amplitude, spike duration (Barthó et al., 2004; Mitchell et al., 2007), ratio of peak to trough (Andermann et al., 2004; Hasenstaub et al., 2005), repolarization slope after trough (Niell and Stryker, 2008), and recovery slope after peak (Figure 2B). The distribution of features from different brain regions is plotted in Figure 2C (Supplementary Figure 2 shows mean and confidence intervals). Previous studies have used these parameters, particularly spike duration, to separate extracellular waveforms into two classes labeled ‘fast spiking’ (FS) and ‘regular spiking’ (RS) (Mitchell et al., 2007; Niell and Stryker, 2008). We observed that in the neocortex and hippocampus, the distributions of spike duration were bimodal, indicating the presence FS and RS subpopulations in these regions (Figure 2C).

**Figure 2.**
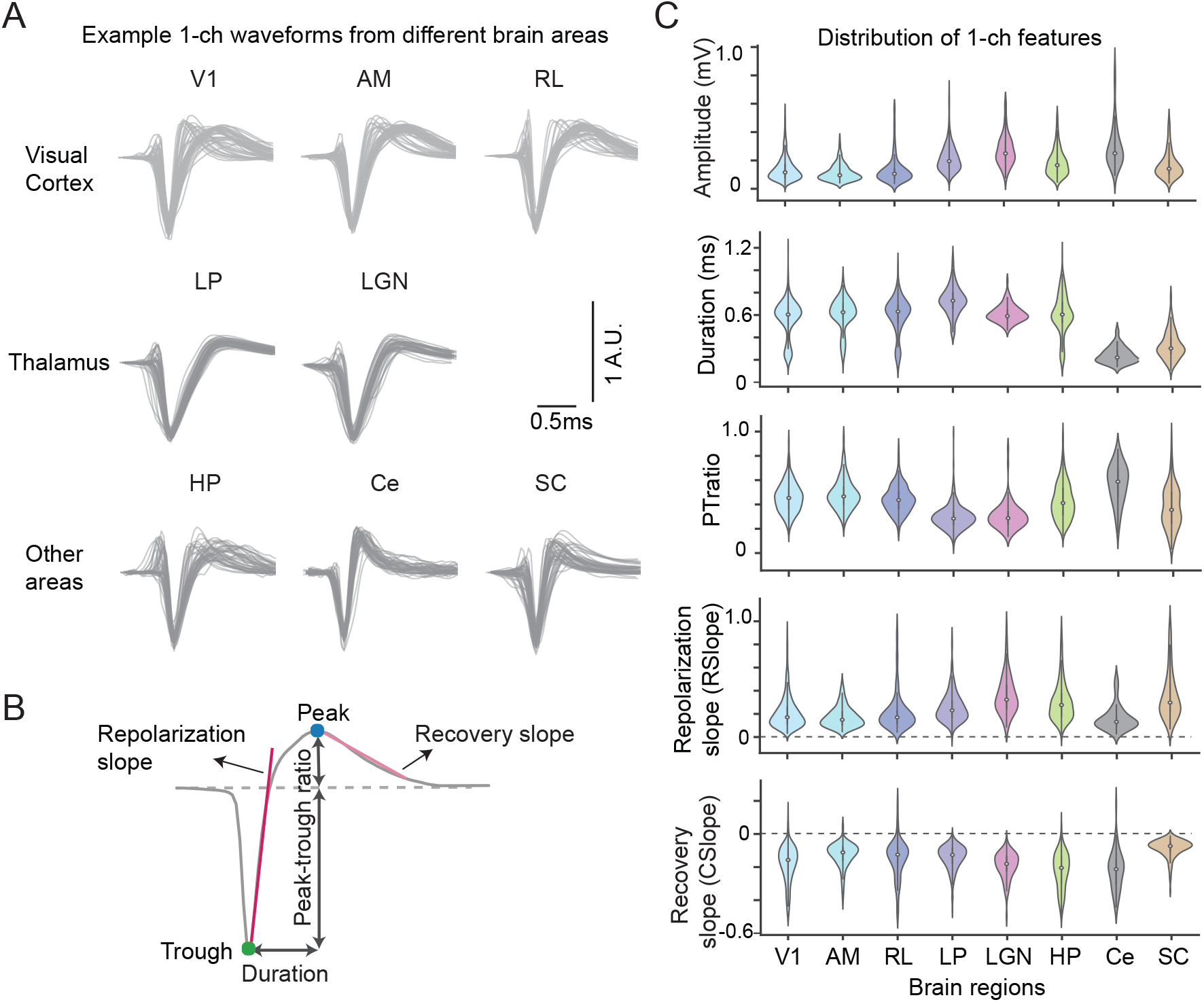
Spike waveform features extracted from 1-channel waveform. **A)** Example normalized mean waveforms for single units from 8 different brain areas (gray lines show 50 randomly sampled single units for each area; waveforms are normalized by amplitude). **B)** Illustration of features extracted from 1-channel waveform (green and blue dots indicate trough and peak, respectively). *Amplitude* is the absolute difference between trough and peak. *Duration* is the time between trough and peak. *Peak-to-trough ratio* (PTratio) is the ratio between amplitudes of peak and trough. Red lines show slopes for repolarization and recovery. **C)** Distributions of 1-channel features in different brain regions. The violin plots show feature distributions estimated with a kernel density function. The white dot indicates median, the thin line shows 95% bootstrapped confidence interval, can colors correspond to different areas (V1, n=1111; AM, n=234; RL, n=264; HP, n=369; LP, n=485; LGN, n=106; SC, n=171; Ce, n=78 units).

In general, we found that 1-channel waveform features are significantly different across brain areas (Figure 2C; ANOVA 1-way test: amplitude F_score=102; duration F _score=190; PT-ratio F _score=131; RSlope F _score=60; CSlope F _score=57, all p_value<<0.001; F-score is used to inform whether a group of variables are jointly significant, and is calculated by variance of group means divided by mean of within group variances). Post hoc comparisons using the paired t-test with Bonferroni correction (see Supplementary Figure 2 for all comparisons) indicates that cortical neurons have smaller spike amplitudes compared to the subcortical neuron types we recorded. The duration and PT-ratio of units recorded in visual cortex and hippocampus are largely similar (p>0.05), but differed from units recorded in the superior colliculus, cerebellum, and LP (p<0.01). Cortical neurons have larger PT-ratio compared to units in thalamus (p<0.05). Units in LP have the longest duration (0.73±0.12ms, standard deviation) and smallest peak-to-trough ratio (0.29±0.09) compared to units from other brain regions. Waveforms recorded from superior colliculus (0.33±0.12ms) and cerebellum (mean duration=0.24±0.07ms) are significantly narrower than other brain regions (p<<0.01).

### Distinct multi-channel waveforms measured in different brain regions

Compared to traditional single electrode and multi-electrode arrays, one advantage of the Neuropixels probe is the relatively dense arrangement of recording sites. Signals from a single unit can be detected on many recording channels (see Figure 3f in (Jun et al., 2017)), providing an additional dimension (space) to characterize cell type-specific spike properties. We defined a multi-channel spike waveform to include the maximum spike channel (closest to soma) and 10 additional channels above and below the peak channel, spanning ±200μm along the probe. This multi-channel spike waveform can be visualized as a heatmap or as a series of voltage traces for each electrode channel (Figure 3A). Figure 3B shows example multi-channel waveforms from eight different brain regions (with additional single unit examples shown in Supplementary Figure 3). The spatial extent of spike waveforms varied across areas. In addition, we noticed that the trough of many spike waveforms appeared to propagate along the linear probe (see visual cortical and HP units in Figure 3A, B and Supplementary Figure 3) (Bereshpolova et al., 2007; Buzsáki and Kandel, 1998).

**Figure 3.**
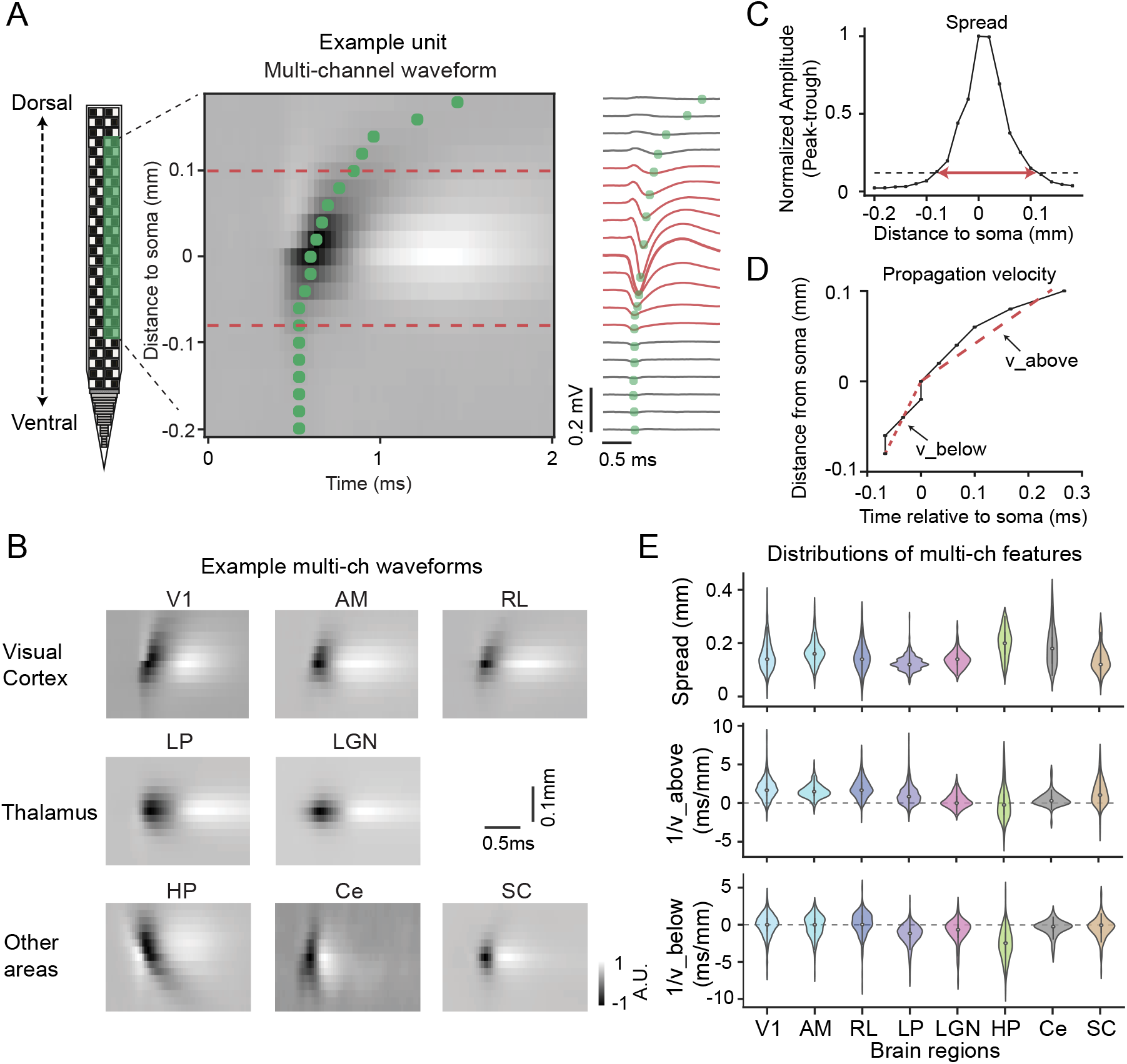
Features extracted from multi-channel waveform. **A)** Illustration of multi-channel extracellular waveform of an example unit. The probe is inserted along the dorsal-ventral axis of the brain and has two parallel columns of recording sites at each depth; for each unit we used waveforms measured on one column of the probe (see Methods). The multi-channel waveform includes the channel with the largest amplitude and 10 channels both above and below. In the heatmap, each row represents the spike from one recording site over time; this same data is plotted as a time-series to the right. Green dots indicate waveform trough at each recording site. Red dashed line indicates the spread of detectable extracellular waveform along the probe, defined in C). **B)** Example multi-channel waveforms from different brain areas showing diverse profiles (see additional example waveforms in Supplementary Figure 3). **C)** Amplitude of the example unit as a function of recording distance to soma. The spread of the waveform along the probe is defined as the distance over which the spike amplitude is larger than 12% of the maximum amplitude. For the example unit, channels within the defined spread are colored in red in A. **D)** Propagation trajectory along the probe for the example unit. For each electrode location (y-axis; y=0 indicates potential soma location) the time of the waveform trough is plotted on the x-axis. Velocity above and below soma are separately estimated by linear regression (red dotted lines). **E)** Distributions of features extracted from multi-channel waveforms in different brain regions (top: spread along probe; middle: inverse of velocity above soma; bottom: inverse of velocity below soma).

To quantify and compare these spatiotemporal spike properties, we next computed several features designed to capture the multi-channel spread and propagation velocity. We define the spread as the distance spanning the contiguous set of electrode sites with spike amplitude larger than 12% of the peak channel (Figure 3C). To characterize spike propagation, we computed the time of the spike trough at each channel within the spread of the spike as a function of channel distance relative to soma (channel with peak amplitude). As shown in Figure 3D the propagation velocity can be computed as the slope of the trough distance versus the trough time. To avoid infinite values caused by the waveform trough occurring on adjacent sites at the same time for some waveforms, we computed the inverse of the velocity separately for the spike propagating above (1/v_above) and below (1/v_below) the cell body location.

The distributions of spread and propagation metrics showed consistent differences across brain regions (Figure 3E; ANOVA 1-way test: spread F_score=82; slope_above F_score=95; slope_below F_score=135, all p_value<<0.001). Post hoc pairwise comparisons using the paired t-test with Bonferroni correction showed that the spread of units in LP, LGN and SC is smaller than other regions (p<0.05; Supplementary Figure 2). Inverse of velocity is indistinguishable among different visual cortical areas (among V1, AM, RL; p>0.05). The inverse of velocity above soma is significantly positive in all visual cortical areas (mean 1.83+0.03 ms/mm; 1 sample t-test p<<0.001), indicating a bias for propagating waves dorsally towards the pia (with mean velocity=0.54 mm/ms). In contrast, in the dorsal hippocampus, where cells are oriented such that the apical dendrites point ventrally, the inverse propagation velocity below soma is significantly negative (−2.60±0.11 ms/mm; 1 sample t-test p<<0.001), indicating a bias for propagating spike waveforms towards the stratum radiatum in CA1 (mean velocity=0.38 mm/ms). Further analysis of propagation profiles will be explored in later sections (Figure 6 and Supplementary Figure 7).

### Brain region classification based on extracellular waveforms

Because extracellular waveforms showed significantly different feature distributions across brain areas (Figures 2-3, Supplementary Figures 2-3), we next tested whether these features could be used to predict which brain region each unit resides in, and whether the multi-channel waveform can provide additional information beyond the 1-channel waveform for classifying unit brain regions. We didn’t include waveform amplitude as a feature for classification since it is strongly affected by the relative distance of the electrode to soma (Gold et al., 2006; Weir et al., 2015). Given the similarity of the waveforms recorded in the three visual cortical areas (VISp, VISam, VISrl), we grouped all the units from visual cortex (VIS) in the following analysis (n=1609 units).

To visualize whether there is any clustering of units in a 2-dimensional space with different waveform feature sets, we first used t-distributed stochastic neighbor embedding (tSNE) (Van Der Maaten, 2008), which is a nonlinear dimensionality reduction technique to embed high-dimensional data in a low-dimensional space for visualization (Figure 4A). Each dot represents one sorted unit and the color indicates the source brain region labeled by histology. Embedding with features extracted from 1-channel waveforms (n_feature=4) is comparatively worse in forming clusters of brain regions; in contrast, features extracted from multi-channel waveforms showed better separation among brain regions with n_feature=7. Using the full single channel waveform (n_feature=60) doesn’t distinguish hippocampus from cortical cells, while using the full multi-channel waveform pushes hippocampus and thalamus far away from other regions. This visualization suggests multi-channel features contain more information to distinguish brain regions than single-channel waveforms.

Next, we trained random forest classifiers to use waveform features to predict the brain region to which each unit belongs. We chose random forest classification because this method minimizes overfitting to training data and makes it possible to assess the contribution of each feature to classification accuracy. To remove potential classification bias that could result from an imbalanced number of units from different brain regions, we subsampled 77 units (without replacement) randomly from each of six brain regions (VIS (including areas VISp, VISam, VISrl), LGN, LP, SC, HP, and Ce). Thus, for a 6-way classification there is a 0.17 chance probability for classifiers trained to predict brain regions. Hyper-parameters for random forest were chosen using grid search with five-fold cross validation (see Methods for details; Supplementary Figure 4). We compared brain region classification based on different sets of waveform features including 1) extracted features (e.g. duration, PT-ratio, inverse propagation velocity), 2) PCA on 1- or multi-channel waveform (90% retained variance), and 3) the entire 1-channel or multi-channel spike waveform (Figure 4B). Classification performance from all feature sets was significantly above chance, with the highest equal to 85.1±1.6% for the multi-channel waveform (standard deviation across 100 different subsamples). Overall, classification performance was improved by using features beyond the traditional 1-channel features, indicating that the spatiotemporal profile of the spike waveforms carries additional information for clustering cells types from different brain regions.

One advantage of random forest classification is its ability to analyze the importance of the features for classification accuracy (Figure 4C). Importance here is defined as the total decrease in node Gini impurity (weighted by the proportion of samples reaching that node) averaged over all the ensembles (Breiman et al., 1984). From our extracted features, the spike duration and PT-ratio were the most important features, and spread was the least important. For data points in the 1-channel waveform, the samples just prior to the trough and around the peak are the most important. For the multi-channel waveform, the samples on the peak channel were important (distance to soma = 0), but there was also a clear contribution of the waveform captured on the spatially adjacent electrode sites within 100-200 um of the soma location, with propagation before and after the trough.

**Figure 4.**
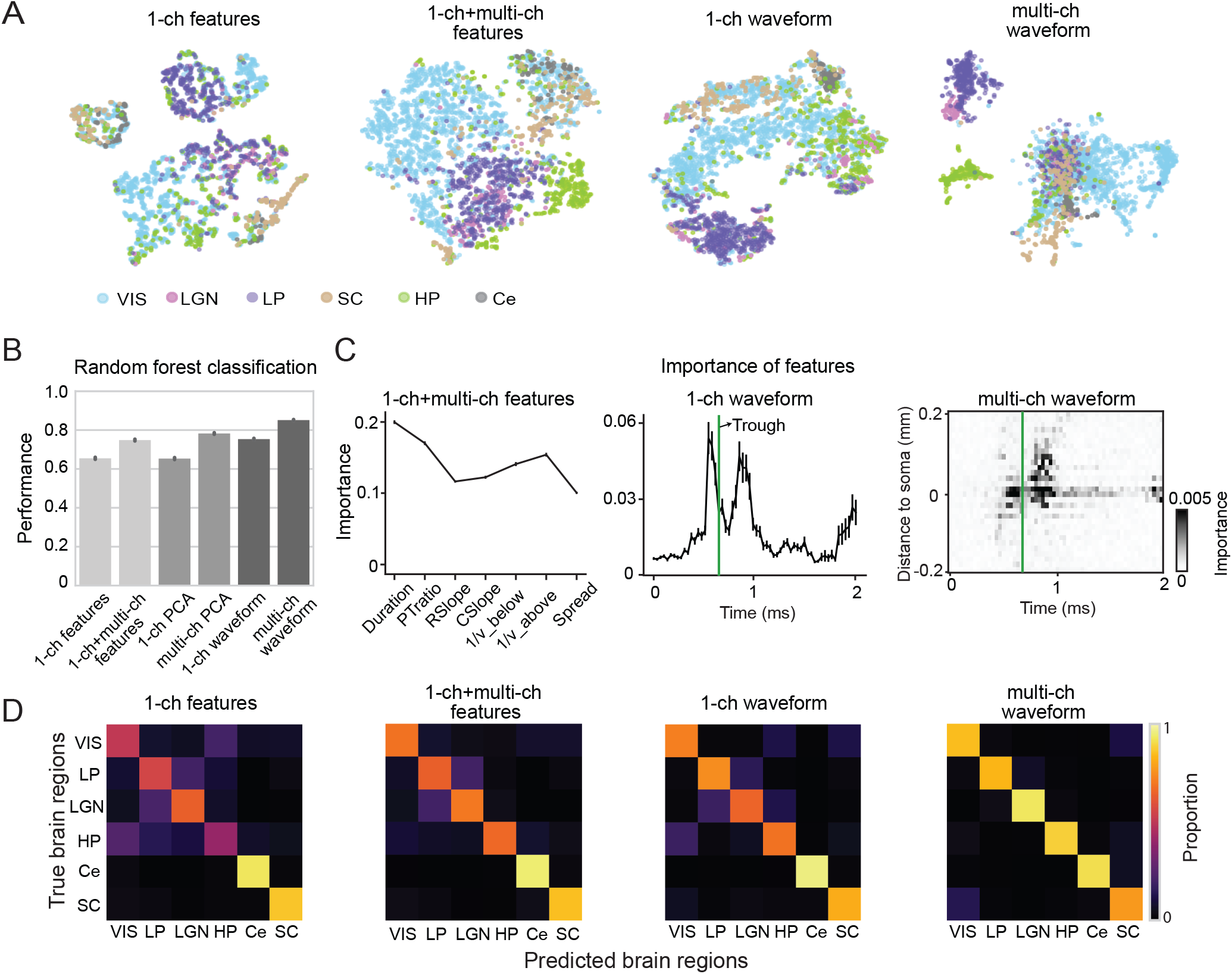
Classification of unit brain region using extracellular waveform features. **A)** t-distributed stochastic neighbor embedding (t-SNE) visualization of units from distinct brain regions based on different waveform features. Each dot corresponds to one unit and brain regions are color coded (n=2818 units). From left, t-SNE representation computed with: features extracted from 1-ch waveform, features extracted from both 1-ch and multi-channel waveforms, 1-ch waveform, and multi-channel waveform. **B)** Performance (out-of-bag score) classifying unit brain region based on 1-channel features alone (n_feature=4), all extracted features (n_features=7), PCA-reduced (90% variance retained) waveform features (1-ch, n_feature=5; multi-ch, n_feature=52), and the whole waveform (1-ch, n_feature=60; multi-ch, n_feature=1800). Error bars are computed over different random samples without replacement (n=100). **C)** Importance of features calculated with random forest classification. Left: importance of extracted features (n_features =7). Middle: importance of 1-ch waveform (n_features =60). Right: importance of multi-ch waveform (n_features =1800). **D)** Confusion matrix of random forest prediction versus true brain regions for different feature sets. Color indicates proportion of units. The average of the diagonals correspond to mean performance in B).

To investigate the misclassification errors that led to imperfect performance with these classifiers, we plotted confusion matrices for different classifiers (with corresponding performance in Figure 4B) to show the predicted versus true brain regions of subsampled units (n=462 units in total, for 77 random samples from each area; Figure 4D). Cerebellum and SC are clearly distinctive from other brain regions for all classifiers. Adding features from the multi-channel waveform can help distinguish hippocampus and thalamus from other brain regions. Interestingly, thalamic units in the LGN could be differentiated from those in the neighboring LP thalamic nucleus using the multi-channel waveform, potentially based on their duration and spread (Figures 2 and 3). Classification without subsampling resulted in higher performance but a similar trend compared across different feature sets (data not shown). Thus, the classification results qualitatively agree with unsupervised embedding, and together suggest that the multi-channel spike waveform profile carries additional information useful for identifying cell classes residing in distinct brain regions.

### Spike waveform clusters within visual cortex

We next examined whether the multi-channel waveform can assist the classification of cell types within a brain region. For this analysis, we focused on waveform types in the visual cortex. The population of visual cortical neurons have a bimodal distribution of spike durations (Figure 2C), suggesting the presence of at least two neuron types (FS and RS). To determine whether multi-channel spike features can identify further waveform types, we applied k-means clustering to the cortical cells. Using the combined 1-channel and multi-channel features, we estimated three waveform clusters (see Methods and Supplementary Figure 5). We visualized these clusters in a 2-dimensional space using a tSNE plot (Figure 5A) with colors representing k-means cluster labels. One of the clusters corresponds to the FS waveform, which includes units with comparatively short duration spikes (Figure 5B). In addition to the FS cluster, which accounted for 19.6% of the units, we identify two regular spiking waveform clusters that we label as RS1 and RS2; these comprise 59.6% and 20.8% of the visual cortex units, respectively. When considering only the 1-channel waveform, the RS1 and RS2 waveforms look very similar in their duration and peak-to-trough ratio (Figure 5B). However, the spatiotemporal structure of the multi-channel spike waveforms is strikingly different between the RS1 and RS2 clusters (Figure 5C). The average RS1 waveform propagates from below the cell body upward along the probe toward the dorsal surface of the brain; in contrast, the average RS2 waveform propagates a shorter distance, symmetrically around soma (Figure 5D). The propagation profile for the FS cluster is also symmetric around the cell body but is relatively flat with a median slope value not different from zero. Overall, the three clusters showed significant differences in several 1-channel and multi-channel features (Figure 5E). FS neurons have significantly shorter duration (ANOVA, p<<0.001) and larger PT-ratio (p<<0.001), with relatively fast bidirectional propagation from soma. Comparing the two RS subclasses: the total spread of RS1 (median=0.16 mm) units is significantly larger than RS2 units (median=0.14; t-test p=9.77E-7); the amplitude of RS1 units (median=0.115) is significantly higher than RS2 units (median=0.098; t-test p=7.67E-7); and the PTratio of RS1 units (median=0.45) is significantly larger than RS2 units (median=0.40; t-test p=1.23E-13). More strikingly, the velocity of waveform propagation above (0.45±0.01 mm/ms; p_value<<0.001 compared to channel-shuffled null distribution) and below the cell body were both positive in RS1 units (1.2±0.02 mm/ms; p_value<<0.001 compared to channel-shuffled null distribution), indicating a unidirectional active spike propagation—likely starting at the spike initiation zone and propagating upward to the cell body and along the apical dendrites of pyramidal neurons. On the contrary, the velocity below soma is negative in FS (−1.2±0.02 mm/ms; p_value<<0.001) and RS2 (−0.7±0.01 mm/ms; p_value<<0.001) units, indicating a bidirectional propagation profile in FS and RS2 cells.

To test whether the waveform clusters we identified via k-means clustering are consistent with known genetically-defined cell types, we performed optotagging experiments in transgenic mice expressing channelrhodopsin in PV+ inhibitory interneurons in the cortex (Pvalb-IRES-Cre; Ai32 (ChR2); Figure 5F). We used a fiber-coupled LED to illuminate the brain surface with both pulse and ramping opto-stimulation patterns to induce a rich response pattern which was used to identify optotagged neurons whose activity profile closely followed the stimulus light pattern (Supplementary Figure 6 and Methods). We optotagged 29 PV+ neurons (4 insertions from two mice) and each of these units is classified as FS based on hierarchical clustering (see Methods). These units clearly fall into the FS cluster shown on the t-SNE plot (Figure 5F; 100% PV+ are FS). The PV+ optotagged neurons have short duration spikes and their multi-channel waveform does not show evidence of unidirectional action potential backpropagation (Figure 5G, H). Thus genetically-identified PV+ visual cortical neurons are of the FS waveform type.

**Figure 5.**
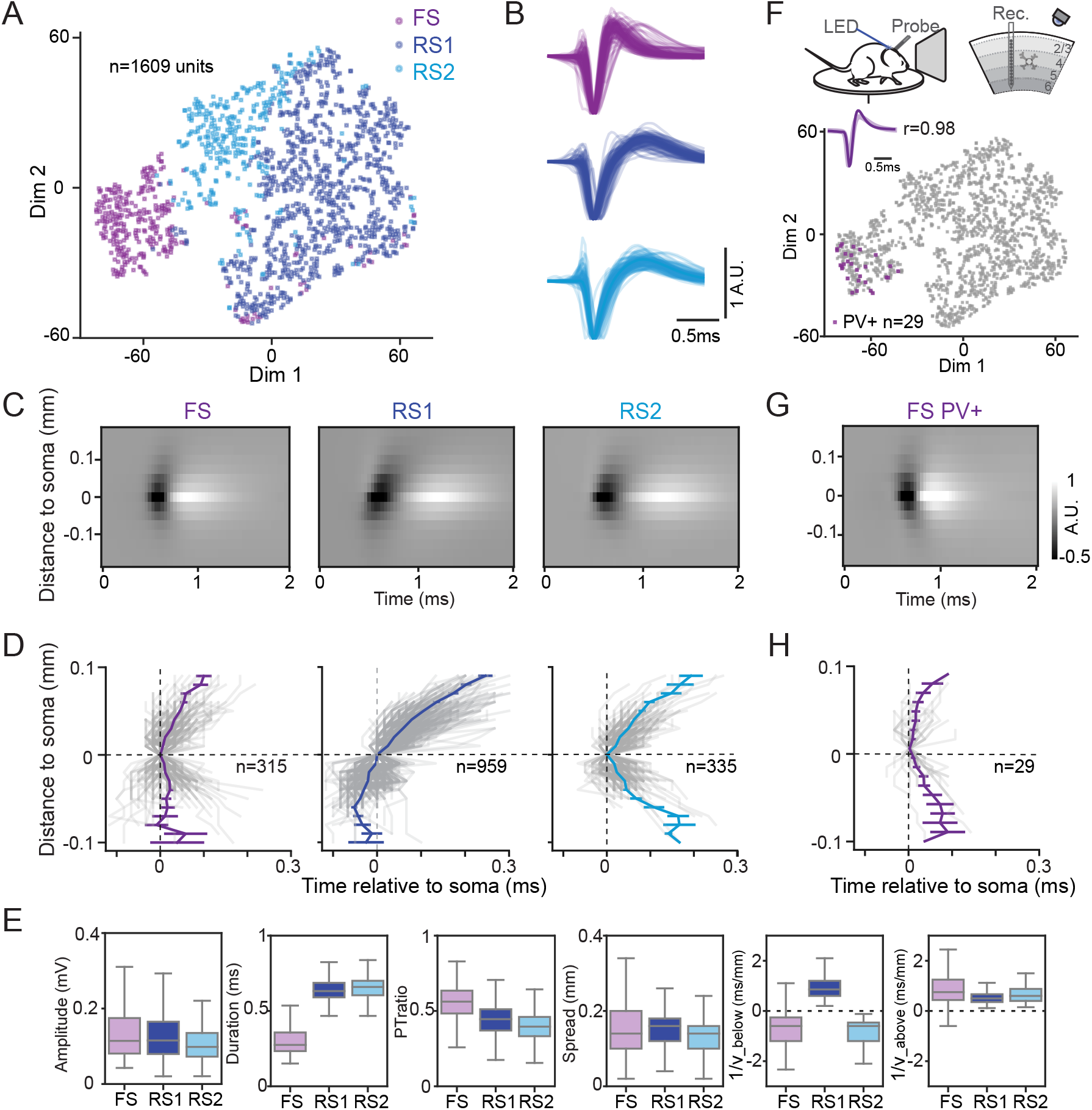
Unsupervised clustering of extracellular waveforms in visual cortex. **A)** t-SNE for all units in visual cortex (n=1609; 11 mice) based on features extracted from 1-channel and multi-channel waveforms. Units are colored according to k-means clustering into 3 groups: fast spiking neurons (FS; n=315); regular spiking neurons type 1 (RS1; n=959); regular spiking neurons type 2 (RS2; n=335). **(B)** Normalized example waveforms for units from different clusters (each trace is the mean waveform of a unit; n=50 sampled units for each cluster). **C)** Average multi-channel waveforms for different clusters. **D)** Spike propagation trajectory from soma for all units in each cluster. Gray lines indicate individual units and the colored lines indicate the mean±sem. **E)** Boxplot of features for different subclasses in visual cortex. The line dividing the box indicates median of distribution, and box represents middle 50% of scores for the group. **F)** Optotagging of PV+ cells in the visual cortex with Pvalb-Cre; Ai32 (ChR2) mice (n=4 insertions from two mice). The top panel is an illustration of the experimental setup for opto-tagging experiments. Probes are inserted vertically in the visual cortex. Blue LED light illuminated the surface of the exposed cortex (peak power=10 mW). Bottom panel is the tSNE representation from A) with color labeled PV+ cells (n=29). The average 1-channel waveform from PV+ cells overlaid on the average 1-ch waveform FS cluster with Pearson correlation equal to 0.98. **G)** Averaged multi-channel waveform for PV+ cells. **H)** Spike propagation trajectories for PV+ cells.

### Backpropagating action potentials in cortex and hippocampus

A previous study showed that backpropagating action potentials in layer 5 pyramidal neurons in rabbit visual cortex are associated with a travelling wave of current sinks and sources along the apical dendrite (Bereshpolova et al., 2007). To determine whether the three waveform types we identified in visual cortex (VIS-FS, VIS-RS1, and VIS-RS2) have distinct patterns of current sinks and sources, we computed the spike-triggered current source density (sCSD; see Methods) profile for each type (Figure 6A). Because sCSD is calculated by second spatial derivative of spike amplitude, it is less sensitive to absolute amplitude and thus the visualization of waveform propagation is more salient. The VIS-FS cluster has a relatively localized current sink centered at the cell body location. In contrast, the VIS-RS1 sCSD profile displays a traveling wave that propagates unidirectionally upward toward the pia (dorsal direction). Interestingly, the electrode sites below the soma have a current sink earlier than the somatic sink—this is consistent with propagation upward from the spike initiation zone toward the cell body (Kole et al., 2008; Stuart et al., 1997). Finally, the sCSD for the VIS-RS2 waveform is also distinct from the RS1 waveform in that the propagation profile is more symmetric around the cell body and the current sinks don’t propagate as far along the probe.

Next, we examined the sCSD for units recorded in the dorsal hippocampus (mostly CA1 region), since backpropagation has also been observed in neurons of this region (Callaway and Ross, 1995; Golding et al., 2001; Jung et al., 1997; Spruston et al., 1995). The hippocampus contains both FS and RS units (Hu et al., 2014), so we first used k-means clustering to divide units into HP-FS and HP-RS types and then computed the sCSD separately for the two waveform types (Figure 6B). The HP-FS cluster did not show evidence of backpropagation, but the HP-RS waveform type displayed a clear sink travelling downward along the probe (ventrally). Thus, the direction of waveform propagation in HP-RS units was the opposite direction compared to VIS-RS1 units. Given the opposite anatomical orientation of dorsal CA1 pyramidal neurons (apical dendrites pointing ventrally) versus visual cortical pyramidal neurons (apical dendrites pointing dorsally toward pia) (Figure 1D), this observation provides further evidence that these propagation events correspond to BAPs along apical dendrites. To visualize the propagation direction for each individual unit we plotted the inverse velocity below versus above the soma (Figure 6C). Most VIS-RS1 units have positive velocities both above and below the soma (p<<0.001 compared to null distribution), indicating unidirectional propagation towards pia. These points fall into the upper right quadrant of the scatter plot. In contrast, many HP-RS units fall into the lower left quadrant indicating negative velocities and unidirectional propagation in the opposite direction.

**Figure 6.**
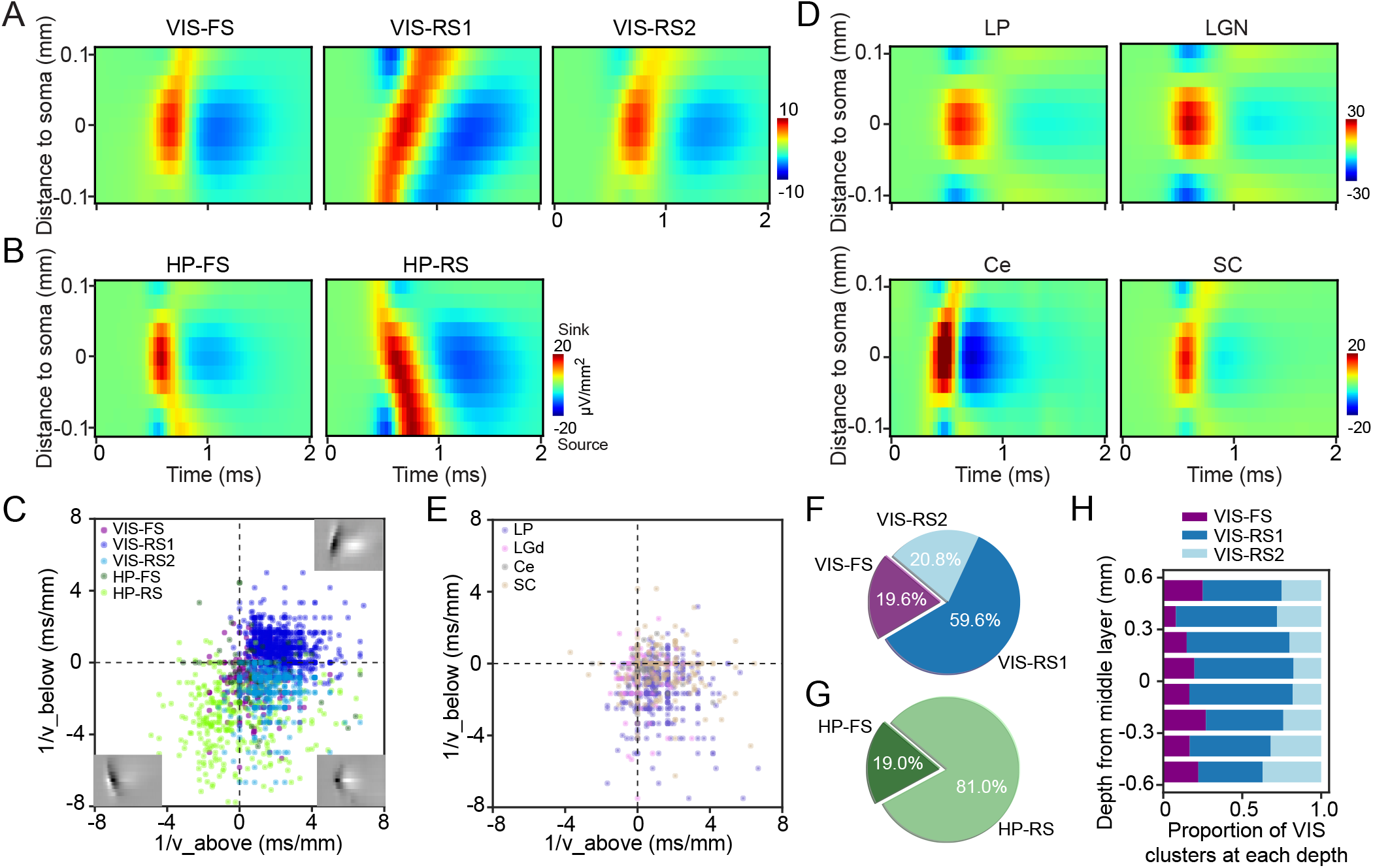
Action potential backpropagation observed in subclasses of cortical and hippocampal units. **A)** Spike-triggered current source density (sCSD) analysis based on the average multi-channel waveforms from different clusters in visual cortex. Propagation towards pia is observed in VIS-RS1 cluster. **B)** sCSD for FS and RS clusters in hippocampus. Note opposite direction of spike propagation in HP-RS compared toVIS-RS1 units. **C)** Scatter plot of inverse propagation velocity (feature extracted from multi-channel waveforms) above and below soma for units in visual cortex and hippocampus (same data as shown in Figure 3E). The upper-right quadrant represents units with unidirectional dorsal propagation and is dominated by VIS-RS1 units. The lower left quadrant represents unidirectional propagation ventrally and is dominated by HP-RS units. The lower right quadrant represents bidirectional propagation from soma, which is dominated by FS and VIS-RS2 units. The multi-channel waveforms (insets) are from example units in three quadrants for intuitive illustration. Values closer to 0 indicate faster propagation speed. **D)** sCSD analysis on the average multi-channel waveforms from LP, LGN, Ce and SC. **E)** Scatter plot of inverse propagation velocity for LP, LGN, Ce and SC. Most of the units are in the lower-right quadrants, indicating the propagation is bidirectional from soma. **F)** Fraction of different subclasses in VIS (n_total=1609 units). **G)** Fraction of different subclasses in HP (n_total=369 units). **H)** Distribution of waveform clusters in visual cortex as a function of cortical depth relative to layer 4. Middle of layer 4 is denoted as 0 and is estimated using stimulus-evoked CSD (see Methods and Supplementary Figure 8).

Importantly, because BAPs are only observed in certain cell types with proper channel composition and morphology, they are known to be absent in some cell types including cerebellar Purkinje cells (Llinás and Sugimori, 1980; Stuart and Häusser, 1994). Thus, we examined the sCSD (Figure 6D) and velocity profile (Figure 6E; Supplementary Figure 7) for subcortical waveforms recorded from the thalamus (LP, LGN), cerebellum, and SC. On average, these regions did not show orientated spike propagation along the probe, with most units occupying the lower right quarter. However, previous findings showed evidence of BAPs in some subtypes of neurons in SC (Gale and Murphy, 2016), which accounts for the observation of some SC units shifted toward the top right of the propagation profile plot (see also Supplementary Figure 7).

Both the VIS-RS1 and HP-RS clusters, which show a high incidence of BAPs, represent a significant fraction of recorded units in visual cortex (56.8%, Figure 6F) and hippocampus (81.0%, Figure 6G). The median velocity of propagation along apical dendrites for VIS-RS1 units is 0.45±0.01 mm/ms (v_above) and the median velocity of propagation along apical dendrites for HP-RS units is 0.30±0.06 mm/ms (v_below). Because the cortex has a layered structure and most previously measured extracellular BAPs events were identified in layer 5 pyramidal neurons in cortex, we evaluated the VIS-RS1 distribution as a function of cortical depth. Interestingly, VIS-RS1 units were observed across the depth of cortex indicating that neurons with BAPs exist in all cortical layers in vivo in the mouse cortex (Figure 6H; also see Supplementary Figure 8). The high spatial resolution of Neuropixels probes (20 μm vertical spacing, compared to 50-100 μm spacing on other linear probes used in previous studies of BAPs (Bereshpolova et al., 2007; Buzsáki and Kandel, 1998)) likely provides the ability to detect events propagating over a shorter distance along the dendrite, and this could be the reason we observe BAP-like events in many regular spiking neurons located in different layers of visual cortex rather than only in layer 5, large pyramidal neurons.

## DISCUSSION

We sought to determine whether detailed analysis of multi-channel spike waveforms captured on high density electrode arrays could assist classification of cell types both within and across different regions of the mouse brain. We measured extracellular action potentials from single units with Neuropixels probes, whose dense recording site arrangement allows detection of extracellular waveforms on multiple probe sites. We found that both 1-channel features such as waveform duration and multi-channel features such as propagation profile were useful for classifying neurons within and between brain regions. However, supervised classification showed significant improvement in performance when operating on the multi-channel compared to 1-channel waveform, indicating that dense spatial sampling of electrical fields helps to detect cell-type specific morpho-electric features. Unsupervised clustering of waveforms in the visual cortex revealed FS and RS units, but also suggests further dividing RS units based on distinct waveform propagation profiles. The propagation profiles of many cortical and hippocampal RS units are strongly indicative of BAPs and demonstrate the potential of the Neuropixels probe for reliable detection of events like BAPs. Finally, we used optotagging to investigate the relationship between our waveform clusters in visual cortex and genetically labeled PV+ inhibitory interneurons. Together, these findings demonstrate the utility of dense extracellular waveforms measured with Neuropixels probes for assisting cell type-specific interrogation of functional circuitry in awake animals.

### Comparison with previous studies

Consistent with previous studies (Barthó et al., 2004; Connors and Kriegstein, 1986; McCormick et al., 1985; Niell and Stryker, 2008; Swadlow, 2003), our work confirmed the separation of FS and RS cells and indicated that PV+ neurons are virtually all FS units (Hu et al., 2014). In addition, there are three main aspects that made our study distinct from previous ones. *First*, the dense sampling of the electrical field by Neuropixels probe allowed us to obtain a rich spatiotemporal profile of the extracellular waveform for each sorted unit, and we demonstrated that this additional information across space can improve cell type classification. The additional features, including spread and propagation velocity, provide additional classification power, and reveal interesting physiological processes such as BAPs that are important for understanding neural computation. Whereas previous in vivo extracellular studies only reported BAPs in cortical layer 5 pyramidal neurons (Bereshpolova et al., 2007; Buzsáki and Kandel, 1998), we detected waveform propagation events in neurons across all layers. *Second*, we compared extracellular waveforms from eight brain regions in this study. Because different brain regions consist of different cell types that express distinct genes and ion channels, we made it an explicit part of our study to compare the diversity of extracellular waveforms from cortical and subcortical brain regions. In the past, most waveform clustering studies have considered spike waveforms only with in the local circuit. *Third*, we applied a diverse set of classification algorithms to analyze cells based on the extracellular action potential. Since we precisely localized each unit to an anatomical region in the brain, we could train supervised random forest classifiers to identify features that are important for cell type classification. In addition, using a combination of low-dimensional embedding techniques and unsupervised k-means clustering algorithms on multi-channel spike features, we identified the presence of at least three waveform types in the visual cortex. Therefore, our study expands the existing knowledge of cell classification based purely on extracellular waveforms.

### Spike waveform and cell types across regions

Our study supports the hypothesis that extracellular waveforms can reflect cell type-specific differences in electrical and morphological properties across brain areas. Multi-channel waveforms were distinct across brain regions, likely reflecting the diversity in morpho-electric properties across areas (Ascoli et al., 2007; Bean, 2007; Stuart et al., 1997; Zeng and Sanes, 2017). Thalamic excitatory neurons typically show a multipolar soma with numerous and highly branched dendrites in a radial or bipolar distribution (Clascá et al., 2012; Jones, 2012). Our results showed symmetric, restricted multi-channel waveforms in LGN and LP neurons, consistent with thalamic relay neuron morphology. Additionally, thalamic relay neurons do not show reliable, long-range dendritic backpropagation (Connelly et al., 2017), which is consistent with our results. Interestingly, we could distinctly classify LGN neurons compared to relay neurons in the adjacent higher order thalamic nucleus, LP. A population of LP relay neurons has been identified with relatively long action potential half-width and afterhyperpolarization potentials (Li et al., 2003); this might account for classification accuracy in these regions of the thalamus and is consistent with the longer spike duration and smaller recovery slope we measured in LP neurons. Our recordings from cerebellar cells also support the view that multi-channel waveforms reflect morpho-electric properties. Purkinje cells in the cerebellum have large cell bodies which can explain the large amplitude and broad spatial spread of the multi-channel waveform. However, because the density of dendritic voltage-gated sodium channels of these cells decreases rapidly with distance from the soma, action potential amplitude drops very quickly in the dendrite and fails to invade the dendritic tree (Llinás and Sugimori, 1980; Stuart and Häusser, 1994; Vetter et al., 2001). Thus, Purkinje cells are known to lack, or have highly attenuated, BAPs (Häusser et al., 2000; Stuart et al., 1997), which is consistent with the lack of obvious BAPs in our extracellular recordings of cerebellar cells. In contrast, many regular spiking units in the cortex and hippocampus showed highly directional dendritic backpropagation suggesting this might be a useful signature for the identification of pyramidal neurons in the cortex and hippocampus.

### Backpropagating action potentials

BAPs are active events that propagate from the spike initiation zone, invade the soma, then travel along dendrites via depolarization of voltage-gated sodium channels or calcium channels (Häusser et al., 2000; Stuart et al., 1997; Waters et al., 2005). BAPs have been studied in vitro and in vivo using both dendritic electrical recording and imaging methodologies (Callaway and Ross, 1995; Jung et al., 1997; Kaiser et al., 2001; Kamondi et al., 1998; Martina et al., 2000; Shai, 2016; Spruston et al., 1995; Stuart et al., 1997; Svoboda et al., 1997; Waters et al., 2005). Two previous studies have used linear multi-channel extracellular electrode arrays to investigate BAPs in the sensory cortex of awake rat (Buzsáki and Kandel, 1998) and rabbit (Bereshpolova et al., 2007). In both cases, the units displaying BAPs were layer 5 regular spiking units; importantly, fast-spiking interneurons did not show these traveling waves. In our study, the enhanced spatial sampling from the Neuropixels probe revealed that many units in the hippocampus and cortex had dendritic propagation events. The velocity of BAPs measured in this study (0.3-0.6 mm/ms) is consistent with the range of velocities previously measured (Bereshpolova et al., 2007; Buzsáki and Kandel, 1998; Shai, 2016; Stuart and Sakmann, 1994). Interestingly, different neuronal types show differences in BAPs, which are caused by a combination of both channel density and dendritic morphology (Häusser et al., 2000; Stuart et al., 1997; Vetter et al., 2001). In pyramidal neurons, BAPs can serve important computational roles including providing a mechanism for synaptic plasticity (Magee and Johnston, 1997; Markram et al., 1997), and supporting dendritic integration of bottom-up and top-down signals (Larkum et al., 1999; Siegel et al., 2000). Moreover, BAPs might be subject to dynamic modulation by behavioral experience (Quirk et al., 2001). Therefore, the ability to routinely measure BAPs in vivo and associate them with functional response properties will be of great physiological importance for understanding the computational roles of spike backpropagation (Häusser et al., 2000; Linden, 1999; Sjöström and Häusser, 2006).

### Influence of probe geometry

One caveat of our study is the influence of probe geometry relative to the morphological layout of individual neurons. Because the waveforms measured by the Neuropixels probe reflect a projection of the extracellular electrical field onto the linear shank of the probe, differences in orientation of individual neurons relative to the probe, and variability of probe insertion angle, can add variability to detected waveform propagation profiles. This is an important factor to consider when interpreting multi-channel spike waveforms. If there is an angle between the probe and apical dendrites of pyramidal neurons, it could cause a reduction in measured speed and spread of BAPs (Bereshpolova et al., 2007). However, for a unidirectional traveling wave, like BAPs along apical dendrites, moderate probe angle variation is not expected to induce a bidirectional propagation profile, suggesting the RS1 and RS2 clusters we identified in visual cortex reflect distinct subpopulations of cells. Biophysical modeling will likely prove important to fully explore how cell type-specific morpho-electric features are reflected in the extracellular spike waveform, and how this depends on factors such probe geometry and electrode sampling density (Buccino et al., 2018).

### Future studies

While our focus here was on single spike waveform features, future studies can include additional features related to spiking pattern and adaptation (Nowak, 2002), spike-train autocorrelation (Ebbesen et al., 2016; English et al., 2017), and cross-correlation analysis to define excitatory and inhibitory cells by inferring monosynaptic interactions (Barthó et al., 2004; Sirota et al., 2008). Combining multi-channel waveforms and spike train features should provide even greater power to reveal distinct cell type-specific properties useful for classification from purely electrophysiological recordings. An important ultimate use of the waveform analysis methods we describe is to study the response properties of different cell classes and their functional roles in complex neural networks. Neuropixels and other high-density probes (Neto et al., 2016; Rios et al., 2016; Scholvin et al., 2016; Shobe et al., 2015) are now being used to generate large-scale datasets in the brain of awake mice performing a variety of sensory, behavioral, and cognitive tasks. Waveform analysis will aid in cell type identification and can also reveal the physiological spike properties such as BAPs that will provide additional insight into the functional logic of neural circuit operations.

## ACKNOWLEDGEMENTS

We thank Ali Williford and Jennifer Luviano for help with surgical procedures, Gregg Heller for help running experiments, Shiella Caldejon for performing intrinsic signal imaging, Timothy Cox for help with tissue clearing and OPT imaging, Will Allen for advice on brain clearing techniques, Marius Pachitariu for help with the spike sorting algorithm. We wish to thank the Allen Institute for Brain Science founder, Paul G. Allen, for his vision, encouragement, and support.

## METHODS

### Data acquisition and preprocessing

#### Data acquisition system

In vivo recordings were performed in awake, head-fixed mice allowed to run freely on a rotating disk. During the recordings the mice either passively viewed visual stimuli or remained in the dark. For recordings in visual cortex (VIS) and hippocampus (HP), data were collected from 11 mice. For recordings in LP, LGN and SC, data were collected from 31 mice (n= 9 in SC, 4 in LGN, and 18 in LP). For recordings in cerebellum (Ce), data were collected from 1 mouse with 3 different penetrations. All spike data were acquired with Neuropixels probes (Jun et al., 2017) with 30 kHz sampling rate, and recorded with the Open Ephys GUI (Siegle et al., 2017). A 300 Hz highpass filter was present in the Neuropixels probe, and another 300 Hz highpass filter was applied offline prior to spike sorting.

#### Animal preparation

For recordings in visual cortex and hippocampus, a metal headframe with a 10 mm circular opening was attached to the skull with Metabond. In the same procedure, a 5 mm diameter craniotomy was drilled over left visual cortex, and sealed with a circular piece of PDMS silicone, ~0.3 mm thick (Heo et al., 2016). Following a 2-week recovery period, a visual area map was obtained through intrinsic signal imaging (Juavinett et al., 2016). On the day of the experiment, the mouse was placed under light isoflurane anesthesia for approximately 40 minutes to remove the silicone window. A ground wire was secured to the skull, and the exposed brain was covered with a layer of 4% agar in ACSF. Following recovery from anesthesia, the mouse was head-fixed on the experimental rig. Three or more Neuropixels probes coated in CM-DiI were independently lowered vertically into visual cortex at a rate of 100 μm/min using a piezo-driven microstage (New Scale Technologies). When the probes reached their final depths of 1200-1500 μm, the tip of each probe extended through visual cortex into hippocampus.

For cerebellar recordings, skin and muscle were resected from above the posterior skull to expose the skull above the cerebellum. Animals were fitted with an aluminum headplate with a 5 mm circular opening above the exposed skull. On the day of recording, the animal was anesthetized and burr holes were made in several locations above the cerebellar cortex. The animal was then head-fixed in the recording apparatus and allowed to recover from anesthesia. For each insertion (n=3 in one mouse), a Neuropixels probe coated in DiI was lowered through a burr hole to a final depth of 3.4 - 3.6 mm from the pia at a fixed rate of 100 μm/min. The probe was fixed at a roughly 15° angle relative to the dorsal plane of the skull, resulting in an 6° to 19° angle relative to the cerebellum surface for each insertion. Recordings extended through multiple ganglionic layers and into the reticular nuclei (Supplementary Figure 1). The probe was allowed to rest in place for at least 15 minutes following insertion before data were recorded.

For LP, LGN, and SC recordings, a metal headframe was attached to the skull with Metabond. One week after surgery, mice were handled (3-5 days) and habituated to head-fixation (~2 weeks). On the day of recording, the animal was anesthetized with isoflurane and a small burr hole (~200 μm diameter) was drilled according to stereotactic coordinates (in mm relative to lambda, LP/LGN: 1.5-2.2 anterior, 1.9-1.5 lateral; SC: 0.25 anterior, 0.5 lateral). Mice were given two hours to recover before being head-fixed in the recording apparatus. A Neuropixels probe was coated in DiI and lowered through the burr hole at a rate of 200 μm/min to a final depth of 3-3.3 (for LP and LGN) or 1.3-1.6 (SC) mm from the brain surface. The probe was allowed to settle for 30 minutes before recording began. For most mice, recordings were made on two consecutive days.

#### Histology

For recordings in visual cortex and hippocampus, the probe location was confirmed by clearing brains with dichloromethane and dibenzylether (https://idisco.info/idisco-protocol/) and imaging with optical projection tomography (OPT; Supplementary Figure 1A). OPT showed most recordings from hippocampus are from CA1 region given our probe insertion location and depth. To assign probes to specific visual areas, we overlaid an image of the brain surface obtained during the recording session on images obtained from intrinsic signal imaging, using the vasculature for registration (Supplementary Figure 1B). For recordings in other brain areas, recording location was subsequently verified by identifying the DiI fluorescence in sectioned brain tissue (Supplementary Figure 1C-F).

#### Data pre-processing

In all experiments, spike times and waveforms were automatically extracted from the raw data using KiloSort (Pachitariu et al., 2016). KiloSort is a spike sorting algorithm developed for electrophysiological data recorded by hundreds of channels simultaneously. It implements an integrated template matching framework for detecting and clustering spikes, rather than clustering based on spike features, which is commonly used by other spike sorting techniques. The outputs of Kilosort were loaded into Phy (Rossant et al., 2016) for manual refinement, which consisted of merging and splitting clusters, as well as marking non-neural clusters as “noise.”

Waveforms for each unit were extracted from the raw data by slicing around the trough time (pre-trough points = 20 samples, total waveform size = 82 samples, with 30 kHz sampling rate). For each unit, the mean waveform was calculated from bootstrapped waveforms (number_of_spikes=100; number_of_repititions=100) from all spikes. If the number of spikes for a given unit was smaller than 100, then all the waveforms were used to calculate the mean waveform. Mean waveforms for experiments with optotagging were calculated only on waveforms prior to the light stimulation period, to avoid artifacts in waveform traces caused by light.

### Optotagging

Optotagging was performed in a subset of the visual cortex experiments described above, using Pvalb-Cre x Ai32 (ChR2 reporter) mice. In each experiment, a 200 micron diameter bare fiber optic cable (Thorlabs) connected to a 465 nm LED (Plexon) was aligned with the center of the cranial window, such that it illuminated a surface area of approximately 80 mm^2^. Stimulus trains were delivered with a Cyclops LED Driver (Newman et al., 2015) and consisted of either 2.5 ms square-wave pulses at 10 Hz, individual square-wave pulses lasting 5 ms or 10 ms, or 1 s raised cosine ramps. Peak light power varied from 0.1 mW to 10 mW on a given trial. Each stimulus condition (pulse type x power) was repeated 120 times. Light artifacts were visible on all channels, but were readily separable from actual spikes based on timing relative to the stimulus and waveform shape.

### Analysis

#### Feature extraction

To classify cell types, we first extracted features from the extracellular waveform. With high density Neuropixels probes, we can record extracellular waveforms of a single unit from multiple sites. We define the recording site with highest amplitude (absolute difference between trough and peak of an extracellular waveform) as the site closest to neuron soma and the extracellular waveform recorded here is our single channel (1-ch) waveform. To take advantage of signals detected by multiple sites, we consider the profile of extracellular waveforms of a single sorted unit recorded from multiple adjacent recording sites as a multi-channel waveform. The probe is inserted along the dorsal-ventral axis of the brain. Since the Neuropixels probe has two recording sites at each depth, we used only the side of the probe with the highest amplitude 1-channel waveforms to calculate the multi-channel waveform. The distance between sites is approximated by their vertical spacing (20 μm). The horizontal spacing is ignored here, but it could potentially contribute to differences between adjacent sites.

For 1-ch waveform, we extracted five features: amplitude, duration, peak-to-trough ratio, repolarization slope, and recovery slope (Figure 2B). Waveform peak is defined by the maximum point of extracellular waveform. Trough is defined by the minimum point. Amplitude is defined by the absolute difference between peak and trough. Duration is defined by the time between waveform trough and peak. This feature is commonly used to separate fast spiking (FS) neurons from regular spiking (RS) neurons (McCormick et al., 1985; Mitchell et al., 2007; Niell and Stryker, 2008; Swadlow, 2003). Peak-to-trough ratio is determined by the absolute amplitude of peak divided by absolute amplitude of trough relative to 0. The repolarization slope is defined by fitting a regression line to the first 30 μs from trough to peak. The recovery slope is defined by fitting a regression line to the first 30 μs from peak to tail.

For multi-channel waveform (Figure 3A), we extracted three additional features in the space domain for classification: spread along probe, inverse of propagation velocity above (1/v_above) and below soma (1/v_below) along the probe. The spread of a unit is defined by the distribution of its waveform amplitude. If we plot amplitude against recording site distance relative to soma, we get a curve with peak at 0 (Figure 3C). We define the range with amplitude above 12% of the maximum amplitude as the spread of a unit along the probe. The multi-channel waveform has information in both time and space dimension for signal propagation velocity estimation. Because the time difference between the trough of adjacent sites could be 0, to avoid infinite numbers, we calculated the inverse of velocity (ms/mm) instead by fitting a regression line to the time of waveform trough at different sites against the distance of the sites relative to soma (Buzsáki and Kandel, 1998).

#### Random forest classification

To classify the brain structure each unit belongs to with extracellular waveforms, we used random forest classification. This supervised learning algorithm provides the contribution of each feature to classification accuracy. In addition, because the accuracy of random forest classification is averaged across many estimators, it is less likely to overfit the data than a decision tree. The two hyperparameters for our random forest classifier, the number of estimators and the depth of the decision trees, were estimated via grid search implemented in Scikit-Learn using 5-fold cross-validation for different feature sets. Because classifier performances for different feature sets plateaued above certain hyperparameter values (Supplemental Figure 4), we chose a fixed set of hyperparameters that reached plateau performance (maximum tree depth=14 and number of estimator = 80) for all feature sets rather than fine-tuning hyperparameters for individual set of features.

All classifications were performed with a 5-fold cross-validation where the classifier is trained on a subset of the data (80%), and then the classifier’s performance is evaluated on the held-out test data (20%). Classification accuracy is determined by the out-of-bag (OOB) score, which is estimated based on the prediction accuracy of data left out in each fit of decision tree (estimator) on bootstrapped sub-samples.

Since we have significantly more units in the visual cortex than other brain regions, we sub-sampled 77 units (determined by n=78 units in cerebellum) randomly from all regions to balance the size of dataset from different brain regions to minimize the influence from underlying class distribution on accuracy. The confusion matrix is computed by comparing predicted classes to true classes for 100 sub-sampled datasets under 100 random initial states. The trend of classification accuracy compared across different feature sets is not different between matched-sample classification and unbalanced-sample classification.

#### K-means clustering

We applied k-means algorithm to determine cell clusters within visual cortex. The k-means method is a widely-used clustering technique that seeks to find centroids that minimize the average Euclidian distance between points in the same cluster to the centroid. However, one of its drawbacks is the requirement for the number of clusters, K, to be specified before the algorithm is applied. We applied two methods in estimating number of clusters in visual cortex (see Supplementary Figure 5). One method is the standard elbow method, which estimates the percentage of variance explained for a given number of K. The number of K is estimated at the point when the curve turns to plateau. Another method estimates the data distribution for a given K, calculated by a density function f(K) (Pham et al., 2005). The value of f(K) is the ratio of the real distortion to the estimated distortion. When the data is uniformly distributed, the value f(K) is 1. When there are areas of concentration in the data distribution, the value of f(K) decreases. Therefore, the number of K cluster is determined by finding the minimum of f(K). Combining estimation of K using the above two methods, we decide on K and apply k-means to data with appropriate number of K for 1000 times with random initial values.

#### t-SNE

t-Distributed Stochastic Neighbor Embedding (t-SNE) is a non-linear dimensionality reduction for the visualization of high-dimensional datasets. We used Laures van der Maaten’s method (Van Der Maaten and Hinton, 2008) to visualize all units in 2-dimentional space with different features. The purpose is to visualize whether units from same structure are mapped to similar regions with a given feature set (Figure 4A). We can also visualize unlabeled data to check whether there are any clusters in units with a given feature set.

#### Classification of opto-tagged neurons

We developed a new method to determine opto-tagged cells based on their response to different light stimulation pattern. Response post-stimulus time histograms (bin size=1 ms) to different light patterns (individual square-wave pulses lasting 5 ms or 10 ms, or 1 s raised cosine ramps) are concatenated to form a response vector for each unit, which result in an n_unit (neurons) by m_feature (time) matrix. We then normalized the matrix and applied PCA to reduce the dimensionality while keeping 90% variance. K-means was applied to the normalized data for 1000 times with random initial value. This process is repeated 100 times to generate a probability matrix with each pixel value indicating the probability of a pair of units belonging to the same cluster. Hierarchical clustering is applied to the probability matrix to determine different clusters. The cluster of units with responses that tightly follow the light pattern is defined as opto-tagged cells. The rest are determined as non-optotagged units. These results were confirmed with labeling based on changes in FR and the significance of those changes (Hangya et al., 2015).

#### CSD

For estimation of unit depth within the cortex, we performed current source density (CSD) analysis by computing the average evoked (stimulus-locked) LFP at each site, smoothing these signals across sites, and then calculating the second spatial derivative (Smith et al., 2013; Stoelzel et al., 2009). For spike triggered CSD, we applied the same metric to the average multi-channel waveform across units (Bereshpolova et al., 2007).

---

**Supplementary Figure 1.**
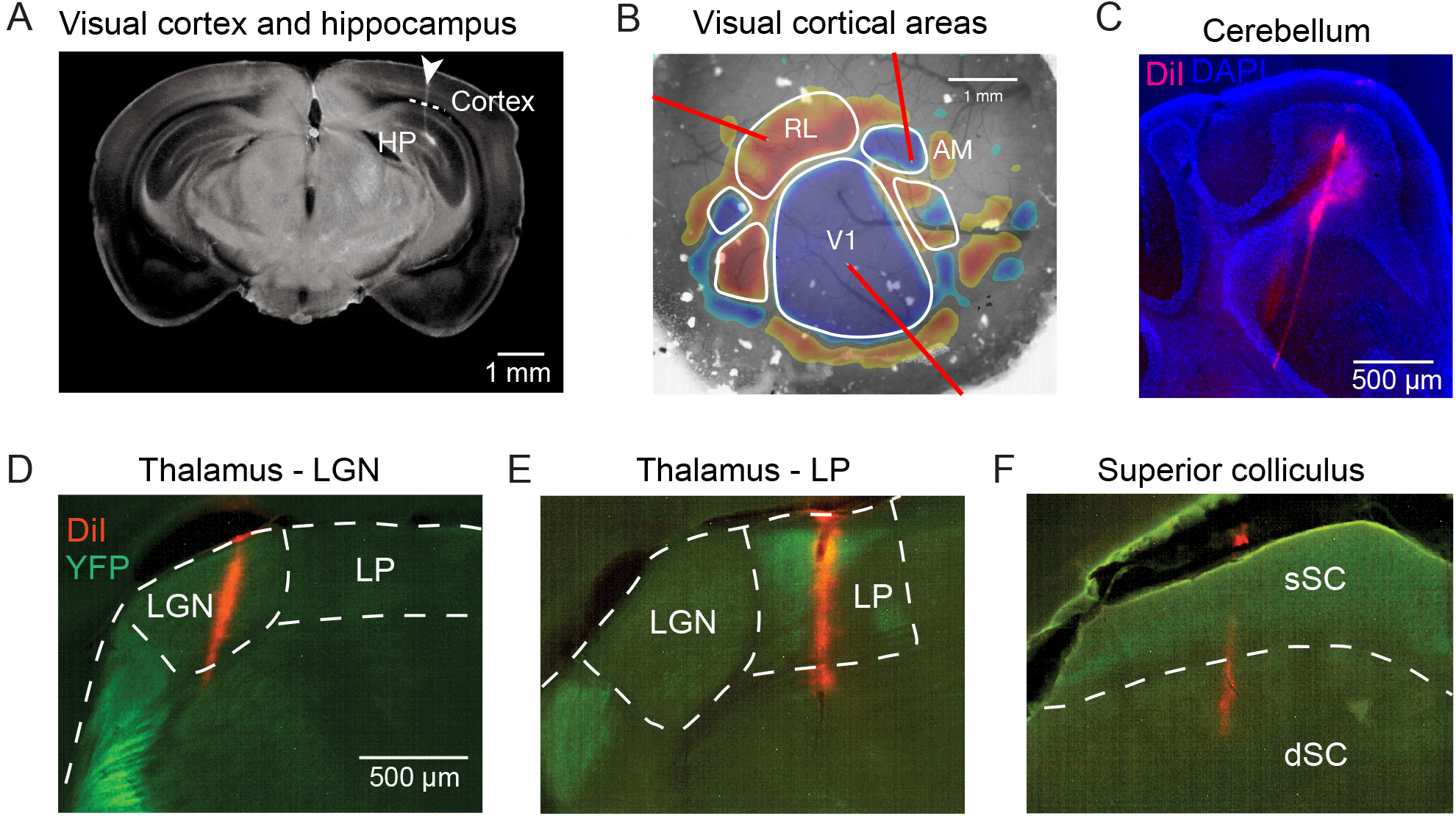
Histological verification of recording location from different brain regions. **A)** Example probe track imaged and reconstructed using optical project tomography (indicated by white arrow). The white matter separates the visual cortex and hippocampus (illustrated with white dashed line). **B)** Overlay of retinotopic sign map and vasculature image to define subareas in visual cortex to guide probe insertion (red lines indicate probes). **C)** Example probe track in cerebellum (red). DAPI (blue) labels cell bodies and shows cytoarchitecture of cerebellum. **D)** Probe track labeled with DiI (red) from example LGN recording in a VGAT-ChR2-EYFP mouse (green labeling shows EYFP in inhibitory neurons of the thalamic reticular nucleus). **E,F)** Probe track verification for LP and SC recordings in NTSR1-GN209 x Ai32 mice. sSC: superficial layers of SC, dSC: deep layers of SC.

**Supplementary Figure 2.**
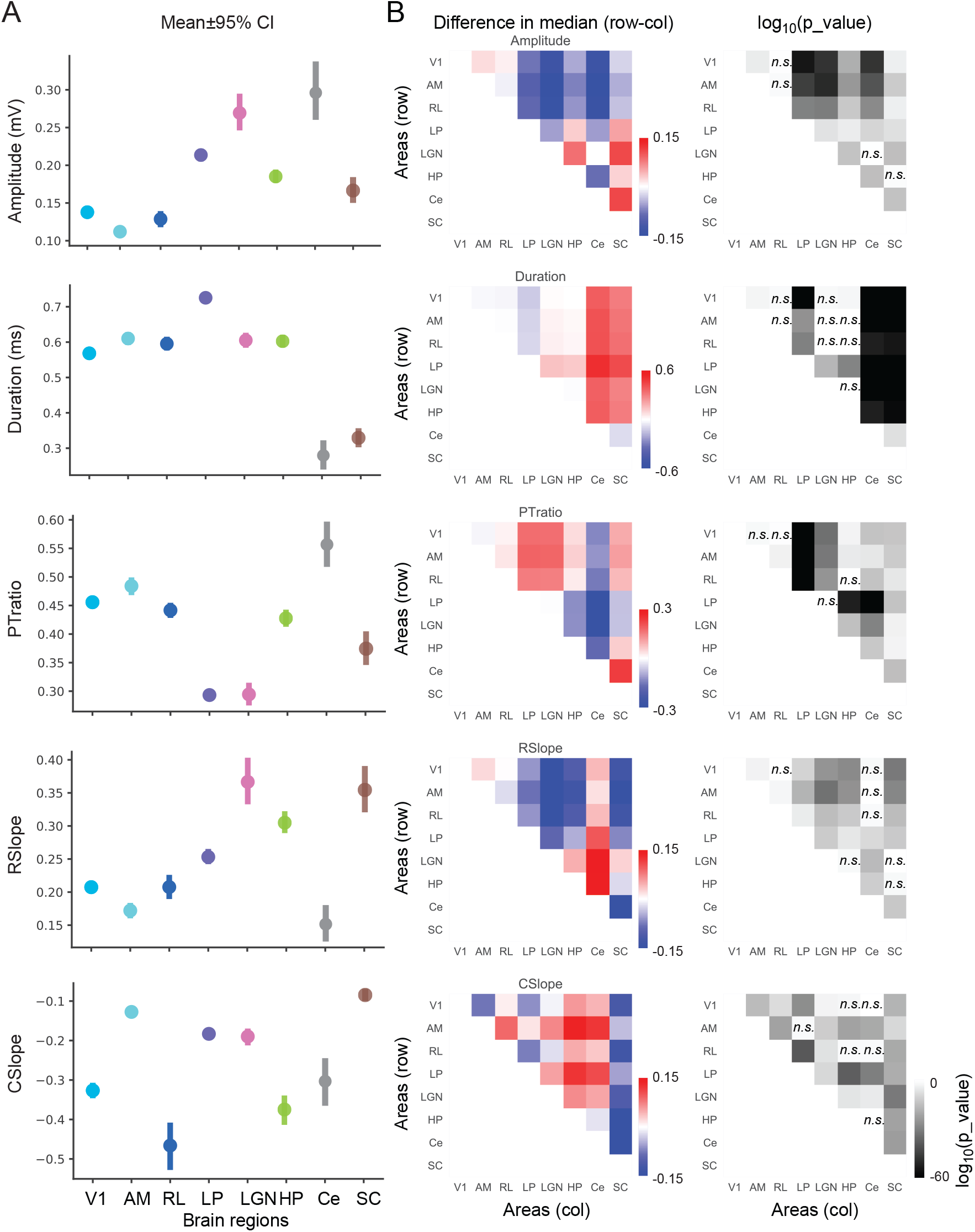
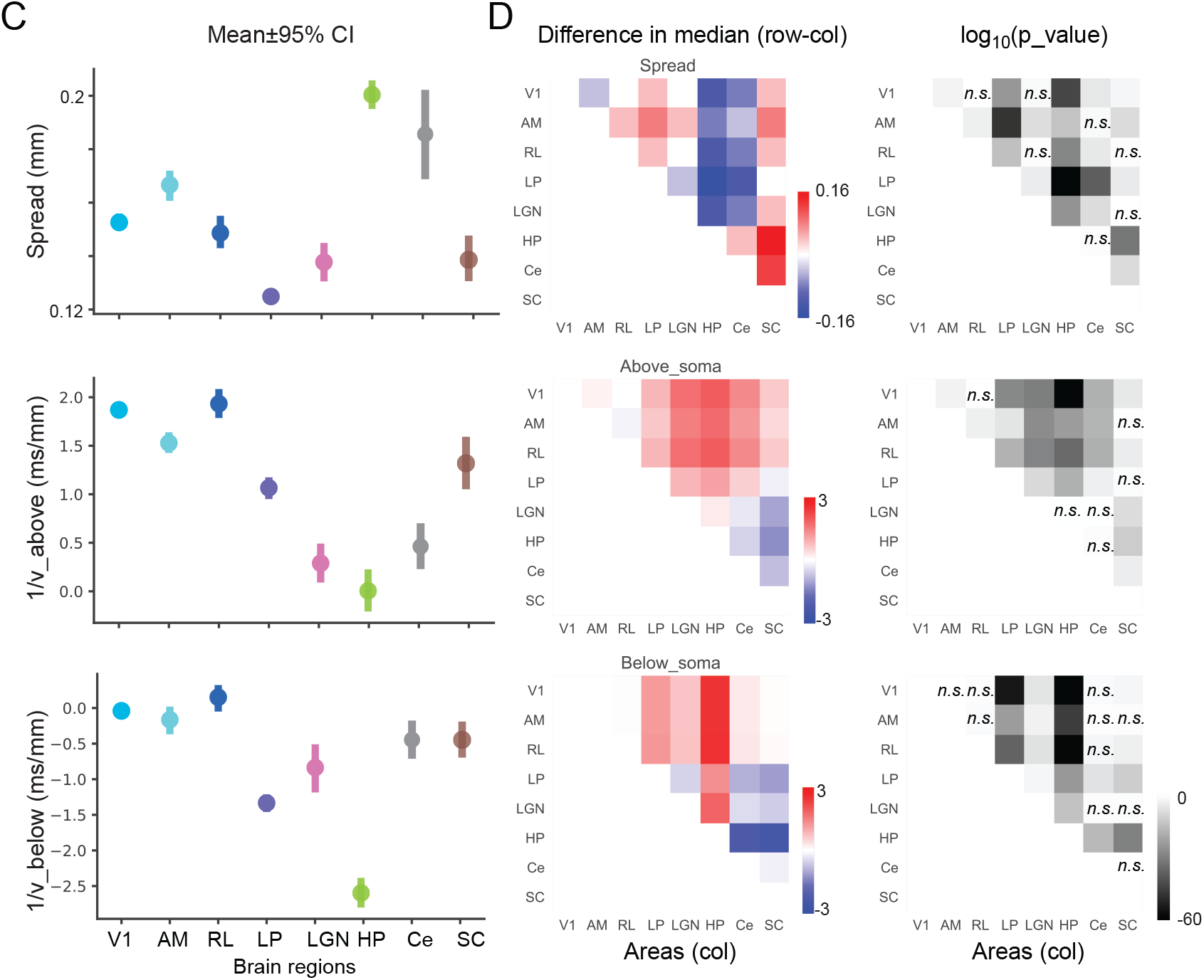
Statistics of extracted waveform features. **A)** Mean and confidence interval (95% confidence interval) of features extracted from 1-ch waveforms from different brain regions. **B)** Pairwise statistics of extracted waveform features. Left panel: Difference of median of the distributions (row-column) for features extracted from 1-channel waveform. Right panel: paired t-test p_value shown in log10 scale, with darker color corresponds to smaller p-value. Significant comparisons were determined by p_value > 0.05 (Bonferroni corrected n=28). Non-significant comparisons are indicated by n.s. **C)** Mean and confidence intervals of features extracted from multi-channel waveforms from different brain areas. **D)** Pairwise comparisons for features extracted from multi-channel waveforms.

**Supplementary Figure 3.**
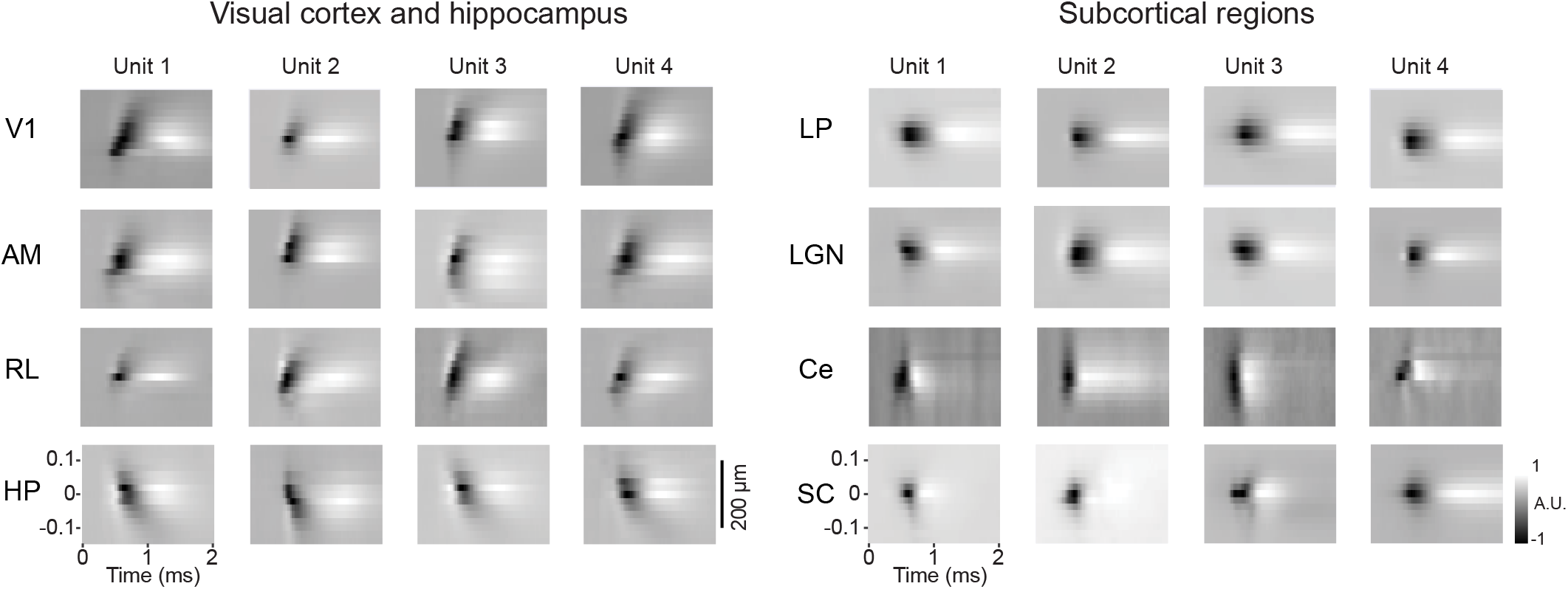
Additional examples of multi-channel waveforms from different brain areas. Four example single unit waveforms from each of eight different brain regions. Each heat map shows amplitude of spike over time, measured on channels above and below the soma.

**Supplementary Figure 4.**
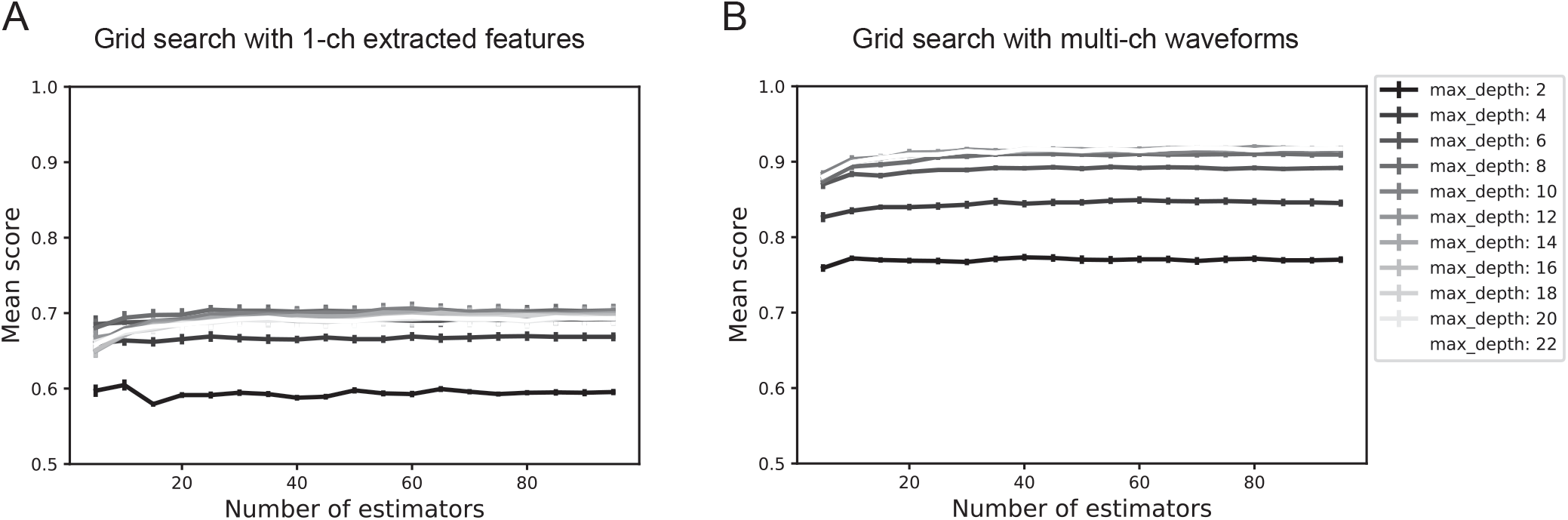
Grid search with different hyperparameters for random forest classification. **A)** 5-fold cross-validation with features extracted from 1-channel waveforms (n_feature=4) as a function of number of estimators (decision trees). Standard error calculated across 5 repetitions. Color intensity of the lines corresponds to different tree depths (see legend). **B)** 5-fold cross-validation with multi-channel waveforms (n_feature=1800) as a function of number of estimators. This result indicates that performance of random forest classification is more sensitive to the maximum tree depth rather than the number of estimators for our dataset. Since performance plateaus above certain tree depth and number of estimators for different feature sets, there is no need to fine tune hyperparameters individually for different feature sets.

**Supplementary Figure 5.**
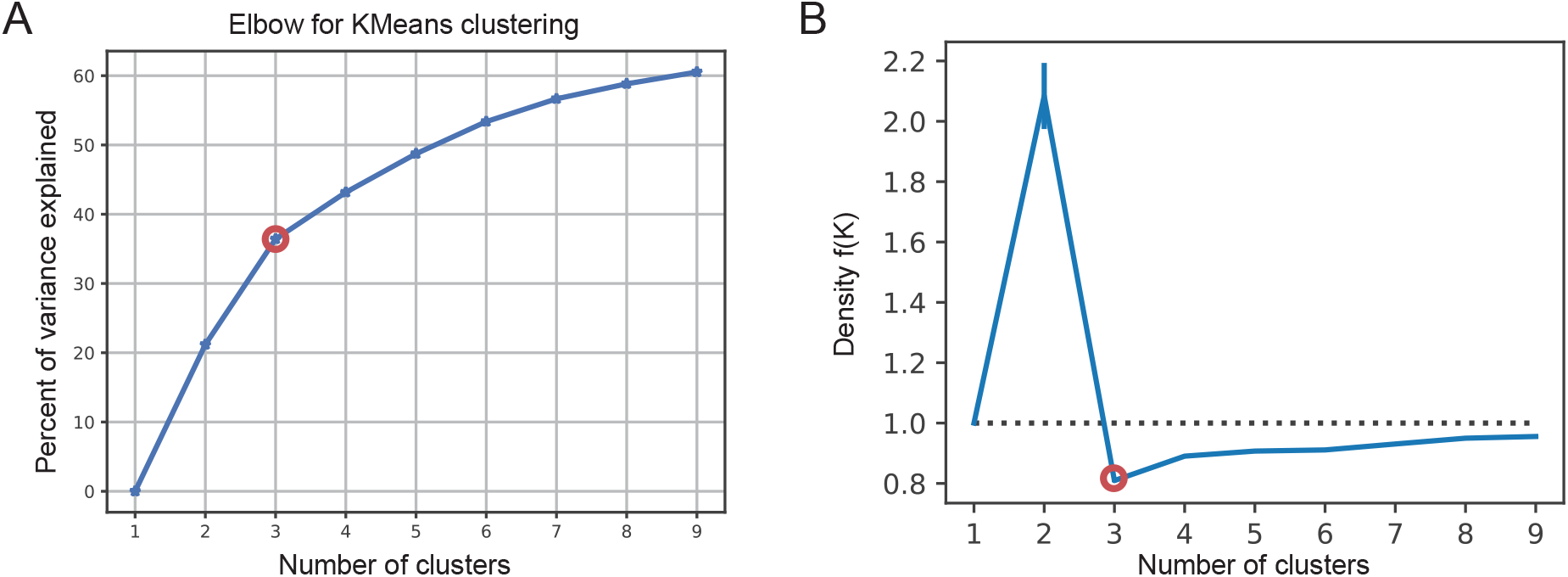
Determining number of K clusters in visual cortex. **A)** Elbow method for finding number of K clusters. Percent of variance explained is calculated as a function of number of K. The turning point is highlighted by red circle. **B)** Density function f(K) as a function of number of k (see Methods). Values lower than 1 indicate concentration of data given the value of K. The number of K clusters that gives rise to the minimum f(K) is determined as the optimal number of clusters (highlighted by a red circle). Both methods indicate K = 3 clusters.

**Supplementary Figure 6.**
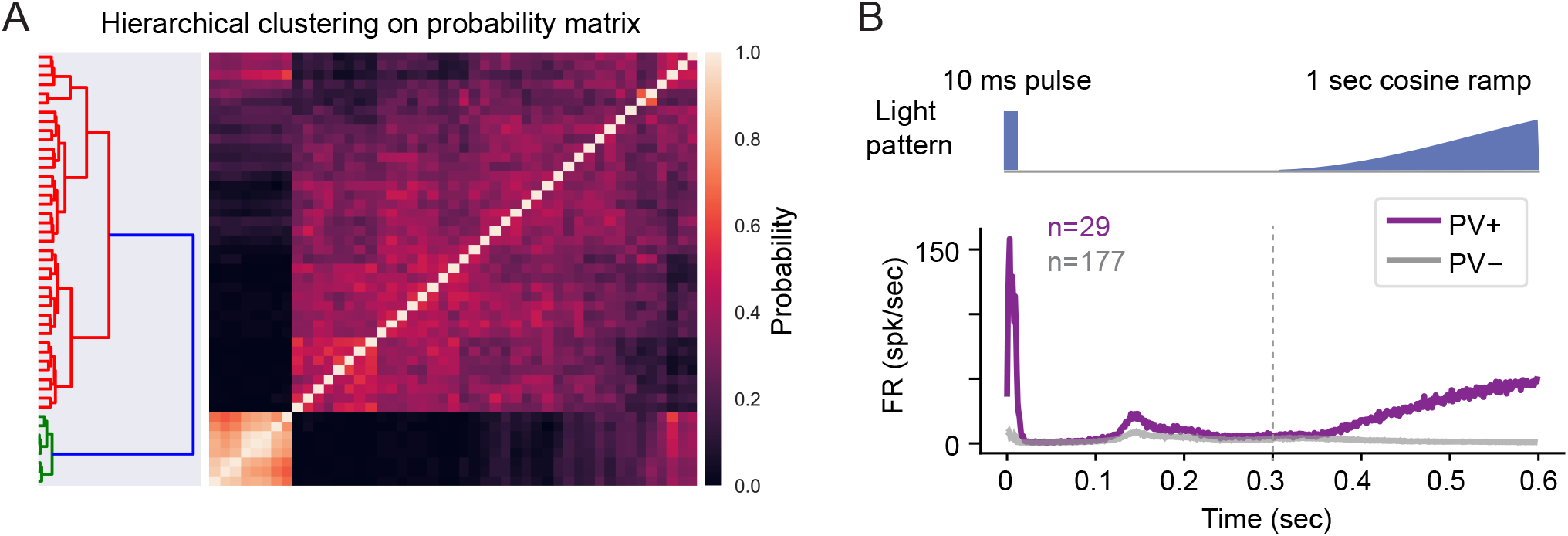
Hierarchical clustering of neuronal response to photostimulation. **A)** Poststimulus time histograms (bin size=1 ms) describing response to photostimulation is used as a feature vector for each unit and K-means clustering is applied 1000 times to the PCA components of the response matrix to construct a co-clustering matrix that reflects the probability of pairs of units belonging to the same cluster. Hierarchical clustering is applied to the probability matrix to find different clusters. This example shows the probability matrix (right) and the hierarchical clustering dendrogram (left) for an example mouse (PV-IRES-Cre; Ai32 (ChR2)) for units in visual cortex. The cluster of units with responses that tightly follow the light pattern is defined as the opto-tagged subpopulation. **B)** Summary plot of mean response to light pattern for all PV+ neurons (n=29 units) from two mice and mean response of the non-opto-tagged neurons (n=177 units).

**Supplementary Figure 7.**
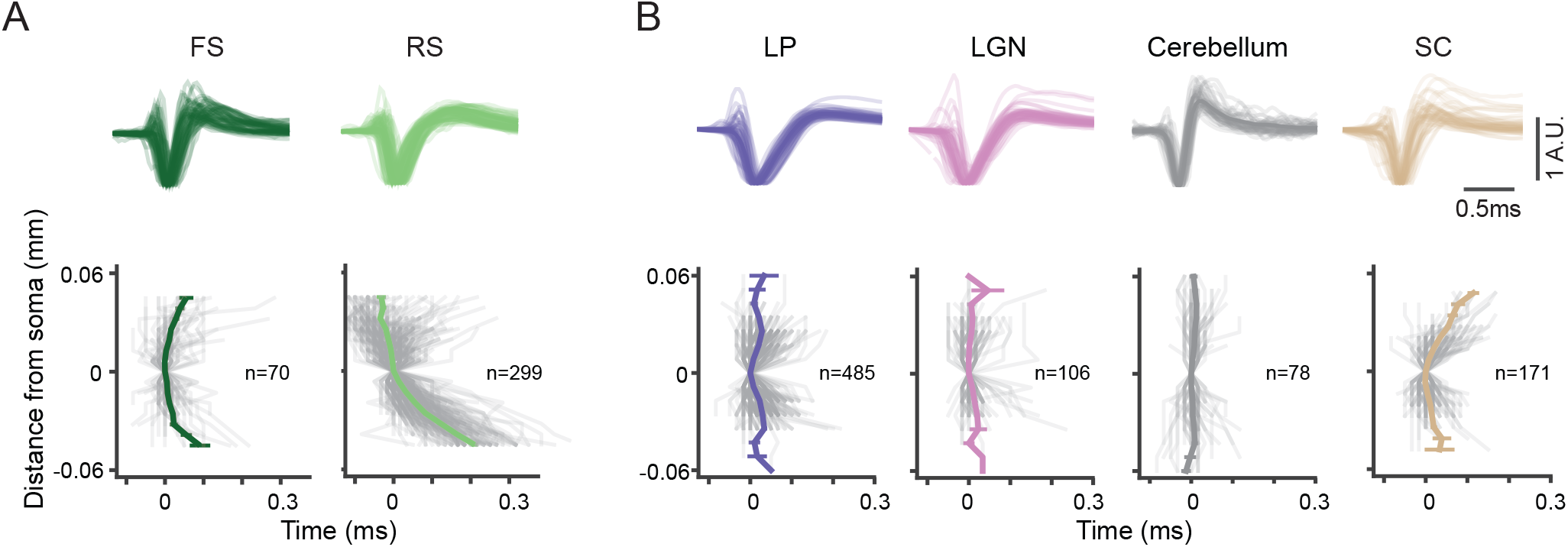
Spike propagation profiles for subcortical regions. **A)** Top: Example waveforms from FS and RS units in hippocampus. Bottom: waveform trough propagation trajectories for all hippocampal FS and RS units. **B)** Top: Example waveforms from LP, LGN, SC and cerebellum. Bottom: propagation trajectories for each region. Gray lines indicate individual units and the colored lines indicate the mean±sem.

**Supplementary Figure 8.**
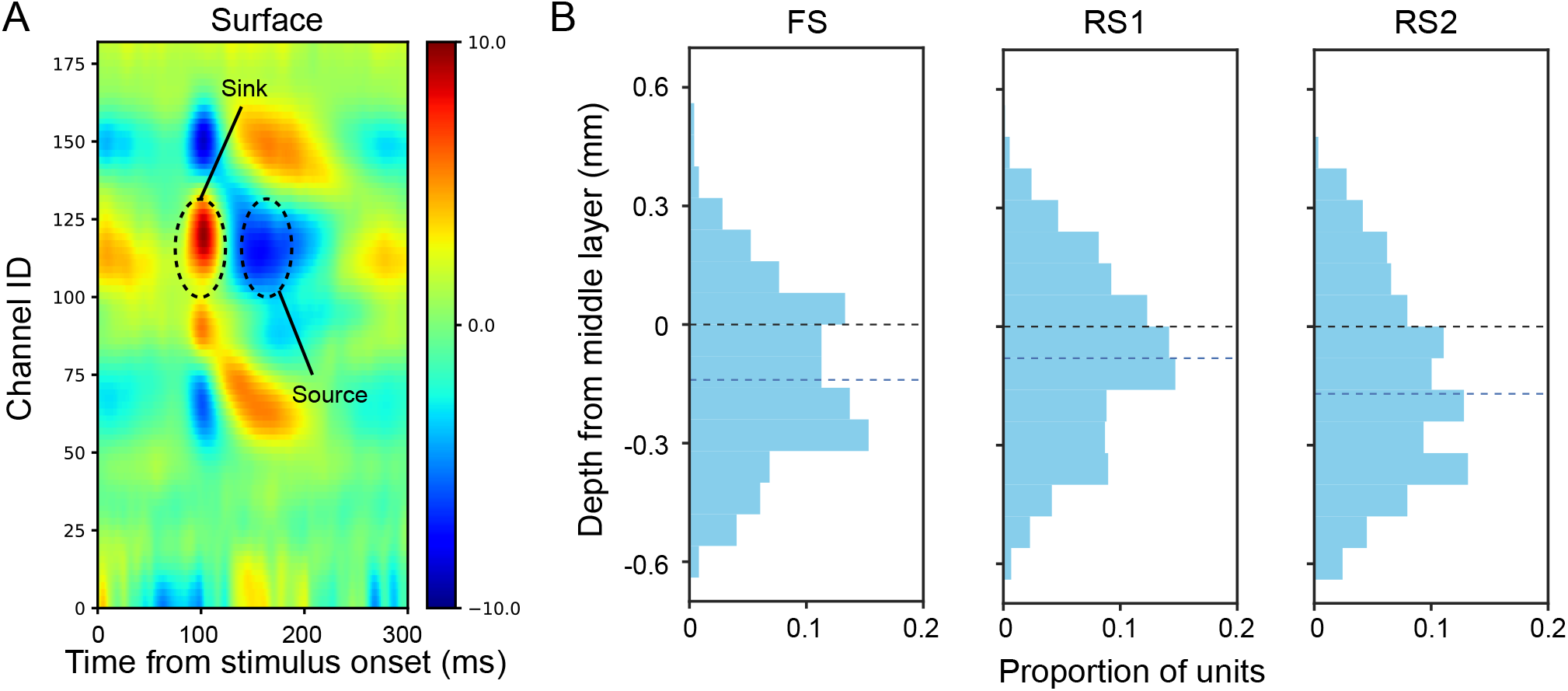
Laminar distribution of visual cortex waveform clusters. A) An example CSD from one insertion. Layer 4 is estimated by the structure of current sinks and sources. Center of mass of the first sink is defined as 0, indicating middle of layer 4. Data from 8 mice (n=1285 units) is aligned to middle layer of visual cortex. B) Proportion of units as a function of laminar depth. Dashed black line indicates depth 0. Dashed blue line indicate median of distribution. The distributions of RS1 and RS2 are significantly different (t-test, p=1.44E-5).

